# Molecular Choreography of Acute Exercise

**DOI:** 10.1101/2020.01.13.905349

**Authors:** Kévin Contrepois, Si Wu, Kegan J Moneghetti, Daniel Hornburg, Sara Ahadi, Ming-Shian Tsai, Ahmed A Metwally, Eric Wei, Brittany Lee-McMullen, Jeniffer V Quijada, Songjie Chen, Jeffrey W Christle, Mathew Ellenberger, Brunilda Balliu, Shalina Taylor, Matthew Durrant, David A Knowles, Hany Choudhry, Melanie Ashland, Amir Bahmani, Brooke Enslen, Myriam Amsallem, Yukari Kobayashi, Monika Avina, Dalia Perelman, Sophia Miryam Schüssler-Fiorenza Rose, Wenyu Zhou, Euan A Ashley, Stephen B Montgomery, Hassan Chaib, Francois Haddad, Michael P. Snyder

## Abstract

Exercise testing is routinely used in clinical practice to assess fitness - a strong predictor of survival - as well as causes of exercise limitations. While these studies often focus on cardiopulmonary response and selected molecular pathways, the dynamic system-wide molecular response to exercise has not been fully characterized. We performed a longitudinal multi-omic profiling of plasma and peripheral blood mononuclear cells including transcriptome, immunome, proteome, metabolome and lipidome in 36 well-characterized volunteers before and after a controlled bout of acute exercise (2, 15, 30 min and 1 hour in recovery). Integrative analysis revealed an orchestrated choreography of biological processes across key tissues. Most of these processes were dampened in insulin resistant participants. Finally, we discovered biological pathways involved in exercise capacity and developed prediction models revealing potential resting blood-based biomarkers of fitness.

## Introduction

Physical activity is a fundamental process associated with improved cardiovascular and immune functions as well as extended lifespan (Ladenvall et al., 2016). In addition to its role in promoting human health, acute exercise is used to identify early signs of pulmonary and cardiovascular disease (Arena et al., 2004). Hence, a detailed molecular characterization of this process is warranted to provide insights into the biological processes involved and the impact of diseases on these processes.

Previous efforts have shown changes in selected circulating markers such as acute inflammatory factors (*i.e.* interleukin-6) (Fischer, 2006) and targeted metabolic pathways (*i.e.* lactate) (Goodwin et al., 2007). However, these studies were limited by the breadth of molecules measured and biological processes covered. In addition, molecular characterization of exercise has mainly been performed in young and healthy individuals (Manaf et al., 2018). Hence, it is not known how metabolic conditions and in particular insulin resistance, affect the response to exercise. Insulin resistance is associated with obesity and type 2 diabetes mellitus which are two major healthcare problems worldwide (Tabak et al., 2012).

In this context, we performed longitudinal multi-omic profiling of blood components (*i.e.* peripheral blood mononuclear cells and plasma) before and after a controlled bout of acute cardiopulmonary exercise (CPX) in 36 participants distributed across a wide insulin resistance spectrum. Symptom-limited CPX testing is a valuable clinical tool that involves acute exercising to maximal oxygen uptake. CPX is used to diagnose lung and heart diseases, assess exercise intolerance and evaluate therapeutic interventions (Arena and Sietsema, 2011; Force et al., 2007). In particular, peak oxygen consumption (peak VO_2_) and the minute ventilation/carbon dioxide production slope (VE/VCO_2_) provide prognostic health information both for the general population as well as for individuals with heart failure and cardiometabolic disease (Arena et al., 2004). Molecules and pathways associated with peak VO_2_ have not been systematically studied, but are expected to be valuable in helping to elucidate and predict human health.

Deep longitudinal molecular profiling revealed an orchestrated choreography of changes involving thousands of molecules in response to acute exercise. Molecular changes were categorized in four dynamic clusters and correlation networks within each cluster illustrated the complex interplay between biological processes across various tissues involved in inflammation, oxidative stress, energy metabolism and tissue repair. We found that myeloperoxidase (MPO) plays a central role in the most acute phase cluster and identified diverse co-occurring molecules including pro-inflammatory (*i.e.* NGAL, IL-7) and growth/protective factors (*i.e.* IL-1ra) as well as complex lipids and acylcarnitines. In addition, we defined healthy molecular profiles of fitness as measured by maximal oxygen consumption, and found important roles for the calpain and integrin pathways in the response to exercise. We also demonstrated the ability of baseline multi-omic analytes to predict important measures of fitness. Finally, we showed that insulin resistance is associated with altered pathways and dampened response to exercise in all major biological processes, and we explored the clinical relevance of molecular outlier analysis for exercise response at an individual level. Altogether, this study illustrates the power of deep longitudinal profiling to unravel the complex physiological response of humans to acute exercise, identify key biological processes involved in fitness and inform impaired mechanisms associated with insulin resistance. Our study also provides a valuable open access resource for the study of the molecular response to exercise.

## Results

### Research design and baseline characteristics of the cohort

Thirty-six highly characterized participants were enrolled and among which 33 agreed to make their data open access (IRB 23602). After overnight fasting, volunteers underwent cardiovascular phenotyping (*i.e.* resting and stress echocardiography and resting vascular ultrasound), symptom-limited CPX testing as well as serial blood collection (**Figure 1a**). CPX testing was performed according to individualized ramp-treadmill protocols allowing participants to reach their maximal exercise capacity within 8 to 12 minutes. Intravenous blood specimens were collected before exercise (baseline) as well as 2, 15, 30 min and 1 hour post-exercise (recovery phase). Additional samples were collected and analyzed at later time points (*i.e.* 2, 4, 6 and 24 hours) but the effect of exercise at these time points was difficult to determine due to the confounding impact of food consumption after the 1- and 4-hour time points. In-depth multi-omic profiling was performed on each sample including gene expression (transcriptomics) from peripheral blood mononuclear cells (PBMCs) as well as targeted and untargeted proteomics, untargeted metabolomics, and semi-targeted lipidomics from plasma. Targeted proteins were selected given their relevance to exercise and consisted of metabolic (n = 9), cardiovascular (n = 39) and immune molecules including 61 cytokines and growth factors (**Table S1**). After data curation and annotation, the final dataset contained a total of 17,662 analytes which included 15,855 transcripts, 260 proteins from the untargeted screen, 109 targeted proteins, 728 metabolites and 710 complex lipids. In this study, complex lipids refer to glycerolipids, glycerophospholipids, sphingolipids and sterol lipids. A list containing all the detected analytes can be found in **Table S2**. The longitudinal multi-omic dataset was used to i) characterize the dynamic molecular response to acute exercise, ii) determine molecular associations with exercise capacity (peak VO_2_) and predict key measurements of fitness, iii) analyze the differential response to exercise in insulin resistant and sensitive participants and iv) examine the clinical relevance of outlier molecules at an individual level (**Figure 1b**).

**Figure 1.**
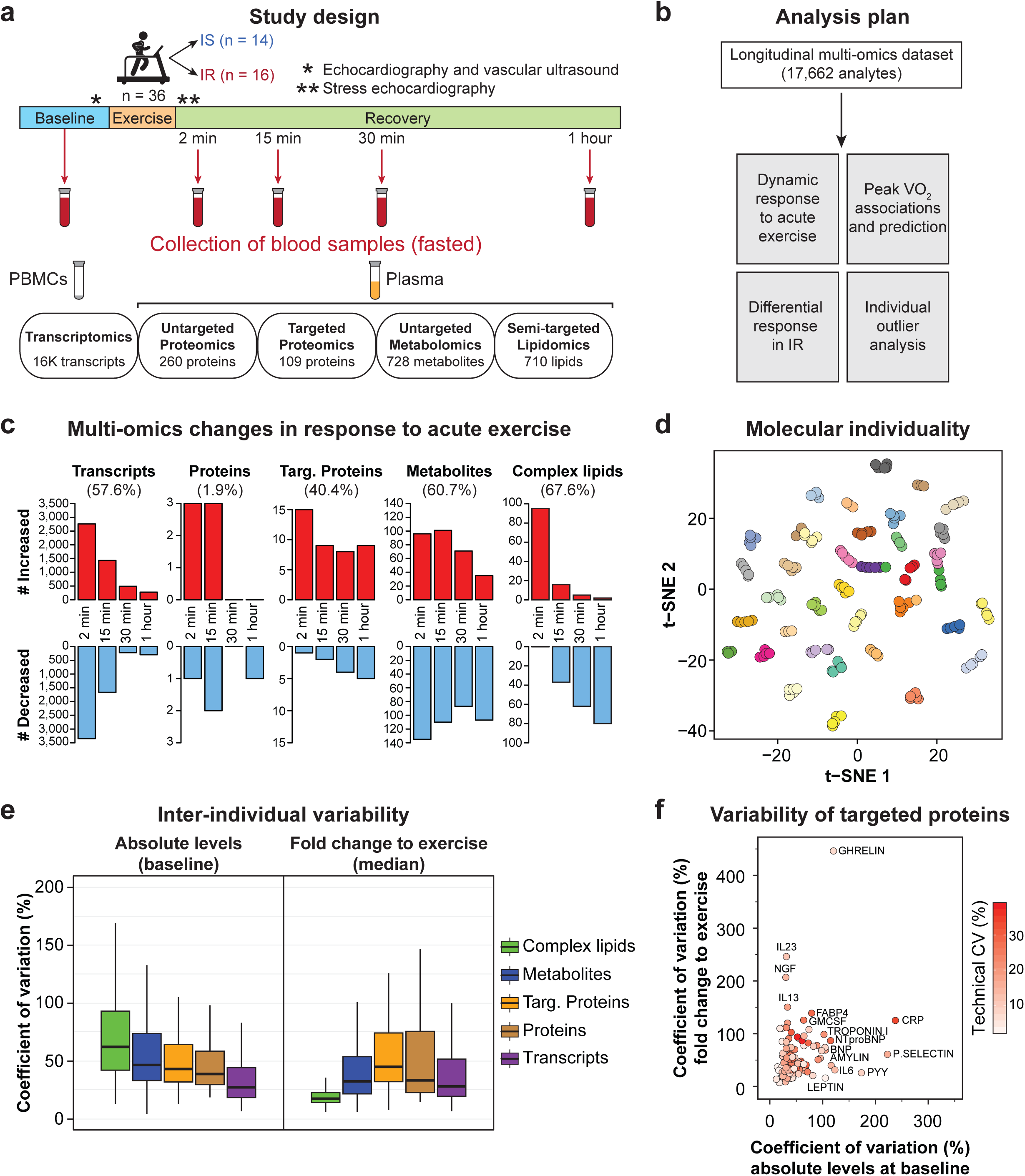
Study design and inter-individual variability. (**a**) Overview of the study design including an acute bout of exercise (symptom-limited cardiopulmonary exercise, CPX), cardiovascular phenotyping and longitudinal multi-omic profiling from blood specimens. IS: insulin sensitive, IR: insulin resistant, PBMCs: peripheral blood mononuclear cells. (**b**) Analysis plan. (**c**) Multi-omic changes in response to acute exercise. Simple linear models correcting for personal baseline, age, sex, body mass index and race/ethnicity were employed. One-way ANOVA testing (two-sided) was then used to calculate significance at each time point. *P* values were corrected for multiple hypothesis using the Benjamini-Hochberg method and analytes with FDR below 0.05 were considered significant. (**d**) 2D visualization of all multi-omic analytes using t-distributed stochastic neighbor embedding (tSNE) technique. Each dot represents a single sample colored by participants. (**e**) Inter-individual variability at baseline (absolute levels) and in response to exercise (delta) across molecule types. The boxplot shows the first (lower edge of the box), median (middle line) and third (upper edge of the box) quartiles. The upper whisker is the third quartile + 1.5 x (interquartile range) and the lower whisker is the first quartile - 1.5 x (interquartile range). (**f**) Inter-individual variability of targeted proteins.

**Table 1.**
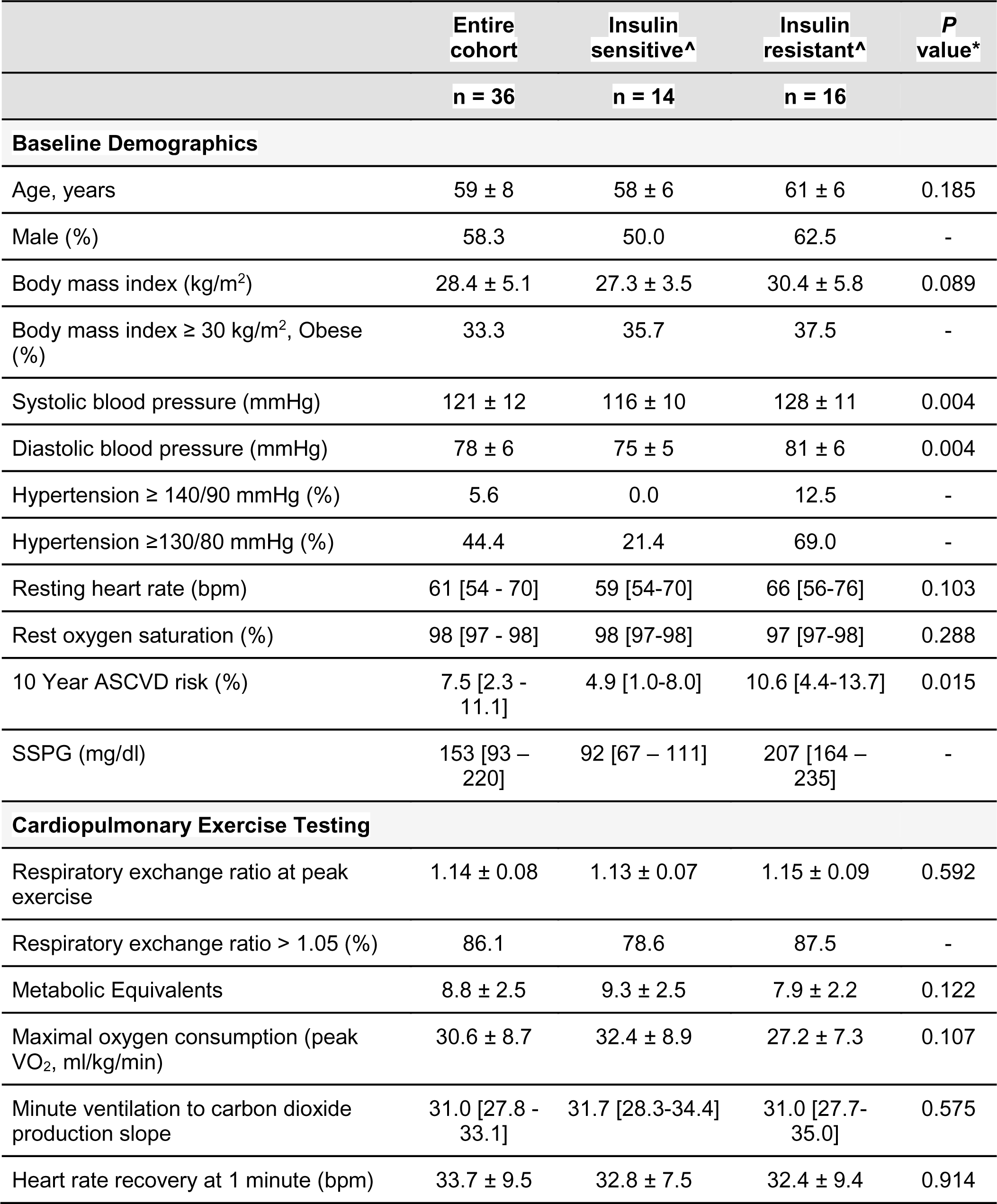
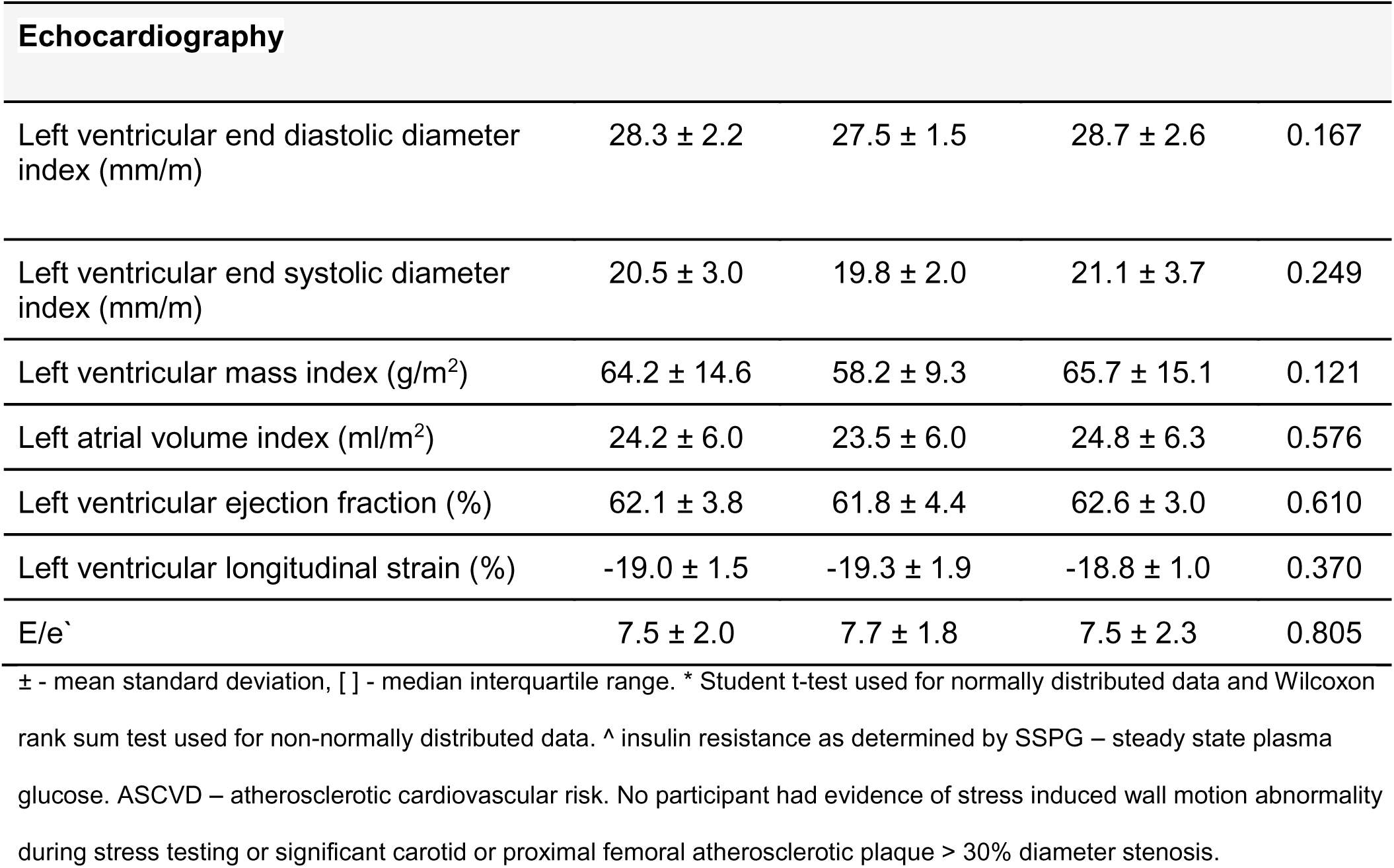
Baseline demographics, echocardiographic and cardiopulmonary exercise characteristics.

The mean age of the cohort was 59 ± 8 years old, the mean body mass index (BMI) was 28.4 ± 5.1 kg.m^-²^ and 58.3% of the subjects were male (**Table 1**). Participants were selected to span a wide range of peripheral insulin resistance with mean steady-state plasma glucose (SSPG) of 153 ± 67 mg/dl as determined by the modified insulin suppression test. Sixteen participants were classified insulin resistant (IR, SSPG ≥ 150 mg/dl) while 14 participants were insulin sensitive. The remaining six individuals did not perform the test because of medical contraindications. The vast majority of participants (86.1%) reached a respiratory exchange ratio (RER) greater than 1.05 at peak exercise and the remaining individuals reached greater than 95% maximal predicted heart rate for their age. The mean peak VO_2_ in the cohort was 30.6 ± 8.7 ml/kg/min, the mean metabolic equivalents (METS) was 8.8 ± 2.5 kcal/kg/hour and the mean minute ventilation to carbon dioxide production slope (VE/VCO_2_) was 31.0 ± 3.8. **Table S3 provides** detailed information regarding participant’s baseline demographics, echocardiographic and cardiopulmonary exercise characteristics.

### Multi-omic changes in response to acute exercise

The quality of each omic dataset was first visually examined by principal component analysis. The study samples were intermixed suggesting limited batch effect and the quality control samples (QCs) clustered together indicating good technical reproducibility (**Figure S1a**). We calculated that 95.2% of metabolites, 91.7% of complex lipids and 65.8% of proteins had a coefficient of variation (CV) below 20% across QCs (**Figure S2a**). One participant underwent CPX testing twice 10 months apart at the beginning and at the end of the study and samples from both sessions clustered together indicating that the exercise protocol and sample collection were reproducible (**Figure S1b**).

Molecules significantly affected by CPX were identified using simple linear models correcting for personal baselines, age, gender, BMI and race/ethnicity. The acute bout of exercise induced extensive changes in 9,815 analytes spanning all omic layers (56.9% of the detected analytes, FDR < 0.05) indicating large system-wide changes (**Figure 1c**, **Table S4**). Different patterns of changes were observed across the molecule types. Transcripts (n = 9,132) exhibited a rapid response reaching a maximum/minimum level immediately post-exercise and returning to baseline (93.7%) within 60 min. In contrast, metabolites (n = 442) and complex lipids (n = 192) were altered across all time points and a large proportion (19.5% and 28.9%, respectively) remained significantly different from baseline 60 min post-exercise (**Figure S1c**). Despite the global molecular impact of exercise, two-dimensional visualization using all multi-omic analytes revealed that samples from the same participant tended to cluster together (**Figure 1d**). Thus, individual molecular profiles were more different than the changes induced by acute exercise.

### Inter-individual baseline versus fold change variability across omic measurements

Molecular profiles are unique to each person; however, it is unknown which classes of molecules and biological processes are the most distinct across subjects at baseline and in response to exercise. Complex lipid profiles varied the most between individuals at baseline (absolute levels) with a median CV of 62.0% followed by metabolites, proteins and transcripts (46.2%, 38.9% and 26.9%, respectively; **Figure 1e**, **Table S5**). A similar pattern was observed at each time point post-exercise (**Figure S2b**). Principal component analysis confirmed these observations with clear individual clusters for complex lipids and metabolites (**Figure S3**). At baseline, triacylglycerol (TAG) and diacylglycerol (DAG) species were the most variable (**Figure S2c**), which is consistent with the clinical variability of total TAG in the cohort (CV = 57.1%). High TAG content variability reflected the wide range of metabolic health status in the cohort from healthy to diabetic. Similarly, xenobiotics – small molecules not produced by the body but typically acquired from the microbiome and the environment – were the most variable among metabolites (**Figure S2d**). For example, a number of metabolites generated in the gut, such as conjugated bile acids (taurocholic acid and tauroursodeoxycholic acid) and indoles (3-Indolepropionic acid), were highly variable between participants. Gene expression data using 412 transcripts with a CV above 100% indicated that inflammation was the most variable biological process in the cohort with a significant enrichment of pathways such as ‘communication between innate and adaptive immune cells’ (FDR = 3.0E-07) (**Table S6**). Differential immune status across participants was confirmed by a number of highly variable proteins including serum amyloid A1 (SAA1) and A2 (SAA2) as well as C-reactive protein (CRP) and interleukin 6 (IL-6) (**Figure 1f**, **Table S5**). These observations validate LC-MS-based proteomic results obtained on a smaller cohort (Geyer et al., 2016).

The pattern of variability in response to exercise (fold change) differed from baseline with proteins varying the most (median CV = 36.8%), followed by metabolites, transcripts, and complex lipids (32.1%, 27.7%, 17.0%, respectively; **Figure 1e**, **Table S7**). The same pattern was observed at each time point taken separately in recovery (**Figure S2e**). Interestingly, 2 out of 21 interleukins (IL-23 and IL-13) as well as NGF (nerve growth factor) presented high variability in response to exercise despite low baseline and technical variability (**Figure 1f**) showing differences among individuals in key immune and nervous system markers. We also observed a different pattern of variability for two key metabolic hormones, acylated ghrelin and leptin, with high and low variability respectively despite similar technical and baseline variability. Leptin is predominantly secreted by adipose tissue whereas ghrelin is primarily produced in the stomach and both play important roles in the regulation of food intake and metabolism (Klok et al., 2007). Both hormones have been reported to decrease following intense exercise and suppress hunger (King et al., 1994). The most variable transcripts in response to exercise (1,334 genes with a CV above 100%) were enriched for ‘osteoarthritis pathway’ (FDR = 5.6E-05) and ‘hepatic fibrosis / hepatic stellate cell activation’ (FDR = 1.6E-04), possibly indicating a direct link between exercise and both osteoarthritis and liver function (**Table S6**).

### Dynamic system-wide molecular response to acute exercise

We took advantage of the high sampling density post-exercise (*i.e.* 2, 15, 30 min and 1 hour) to i) define clusters of longitudinal trajectories using significantly changing plasma multi-omic measurements (FDR < 0.05) and ii) calculate pairwise correlations between molecules within each cluster. Using c-means clustering, four main clusters of longitudinal trajectories were defined (**Figure 2a**, **Table S4**). Some molecules accumulated during exercise and quickly returned to baseline during the recovery phase (cluster 1) whereas others presented a delayed increase post-exercise before returning to baseline (cluster 2). The remaining analytes decreased in response to exercise with some returning to baseline within 1 hour (cluster 3) and others continuing to decrease in recovery (cluster 4). Correlation networks were generated for each cluster and revealed a comprehensive and orchestrated suite of dynamic molecular changes delineating known and novel biological functions associated with exercise. In addition, network structures highlighted key regulators and new potential functions through unexpected connections.

**Figure 2.**
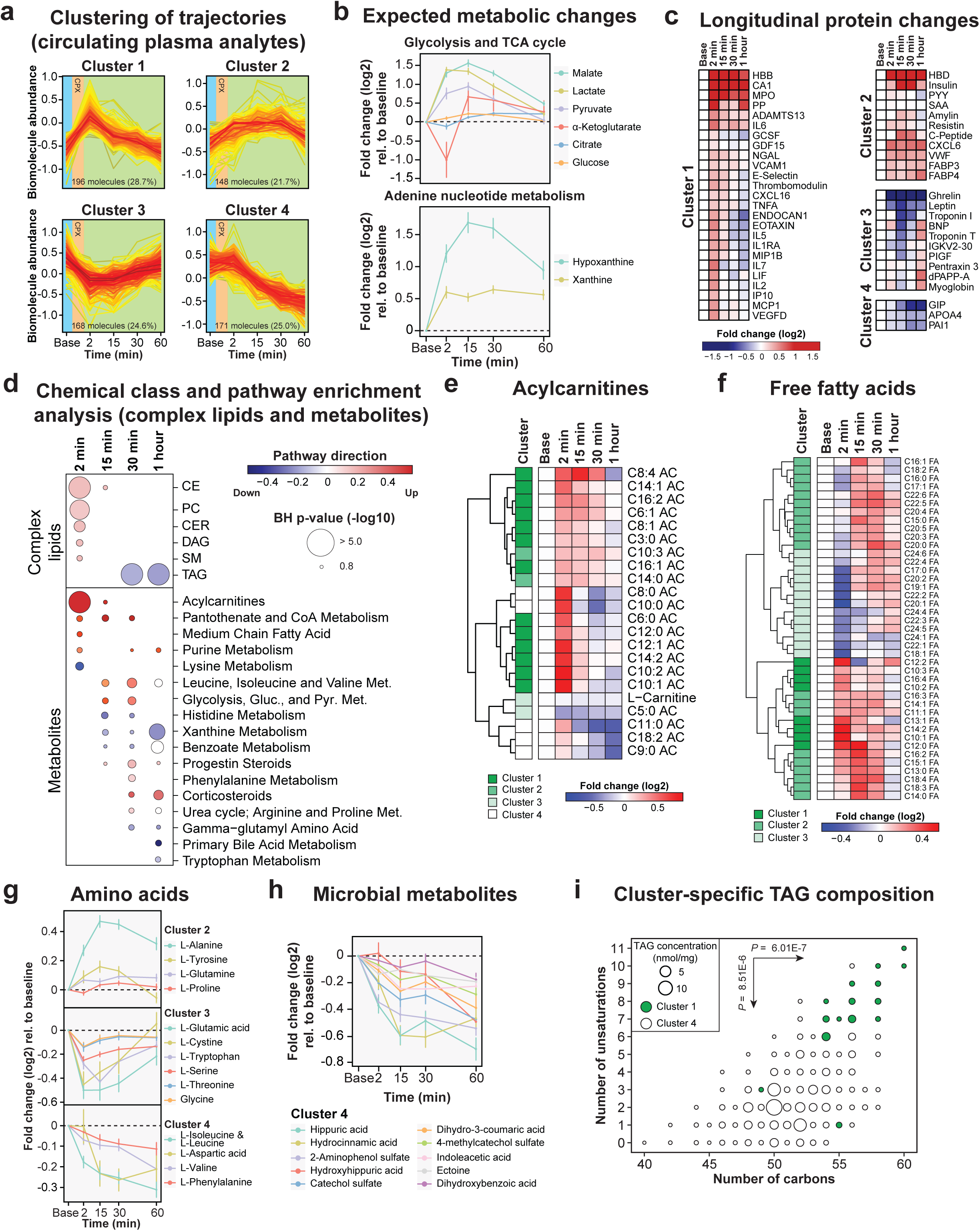
Dynamic system-wide molecular response to acute exercise. (**a**) Clustering of longitudinal trajectories using significant circulating plasma analytes (FDR < 0.05). (**b**) Expected metabolic changes in response to exercise including glycolysis and TCA cycle and adenine nucleotide metabolism. The dots represent the mean log2 fold change relative to baseline and the bars the standard error of the mean (mean ± sem). (**c**) Heatmap of significant proteins representing the median log2 fold change relative to baseline in the cohort. Proteins were grouped by clusters. (**d**) Pathway/chemical class enrichment analysis of circulating plasma metabolites and complex lipids using the Kolmogorov–Smirnov method (one-sided). *P* values were corrected for multiple hypothesis using the Benjamini-Hochberg method and pathways with FDR below 0.20 were considered significant. Pathway direction is the median log2 fold change relative to baseline of significant molecules in each pathway (blue: downregulated, red: upregulated). The dot size represents pathway significance. Heatmaps representing the median log2 fold change relative to baseline for acylcarnitines (**e**) and free fatty acids (**f**). The clusters are indicated on the left side of the heatmaps. Longitudinal trajectories of significant amino acids (**g**) and microbial metabolites (**h**) in response to exercise. The dots represent the mean log2 fold change relative to baseline and the bars the standard error of the mean (mean ± sem). (**i**) Triacylglycerol (TAG) fatty acid composition in clusters 1 and 4. A two-sided Welsh’s t-test was used to calculate differential enrichment in TAG composition including the number of unsaturations and the number of carbons.

#### Cluster 1

Cluster 1 was enriched in molecules (n = 196) associated with anaerobic metabolism, immune response, oxidative stress, fatty acid oxidation (FAO) and complex lipid metabolism (**Figure S4**). We observed a sharp increase in plasma concentrations of glycolysis products (*i.e.* lactate, pyruvate) and tricarboxylic acid (TCA) cycle intermediates (*i.e.* malate) reflecting heightened anaerobic metabolism and TCA cycle flux (**Figure 2b**) (Lewis et al., 2010). These changes are expected and validate the exercise protocol, sample collection and data generation.

We also detected an inflammatory response post-exercise through the secretion of interleukin-6 (IL-6) and tumor necrosis factor alpha (TNF-α) (Golbidi and Laher, 2014) (**Figure 2c**). Pro-inflammatory properties of TNF-α was confirmed by its association with many other cytokines (*i.e.* IL-2, IL-6, IL-7, granulocyte-colony stimulating factor (G-CSF), monocyte chemoattractant protein 1 (MCP-1), leukemia inhibitory factor (LIF) and eotaxin) (**Figure 3a**). Interestingly, TNF-α also correlated with pantothenic acid (vitamin B5) which is required for the biosynthesis of coenzyme A and in turn regulates cellular energy metabolism including fatty acids, carbohydrates and amino acids (Tahiliani and Beinlich, 1991). This observation suggests that in addition to its well-known role in inflammation, TNF-α regulates energy metabolism in the context of exercise and is consistent with a large body of literature (Chen et al., 2009; Sethi and Hotamisligil, 1999). A compensatory anti-inflammatory response to re-establish homeostasis was concomitant with an increase of IL-1 receptor antagonist (IL-1ra) and vascular endothelial growth factor D (VEGF-D).

**Figure 3.**
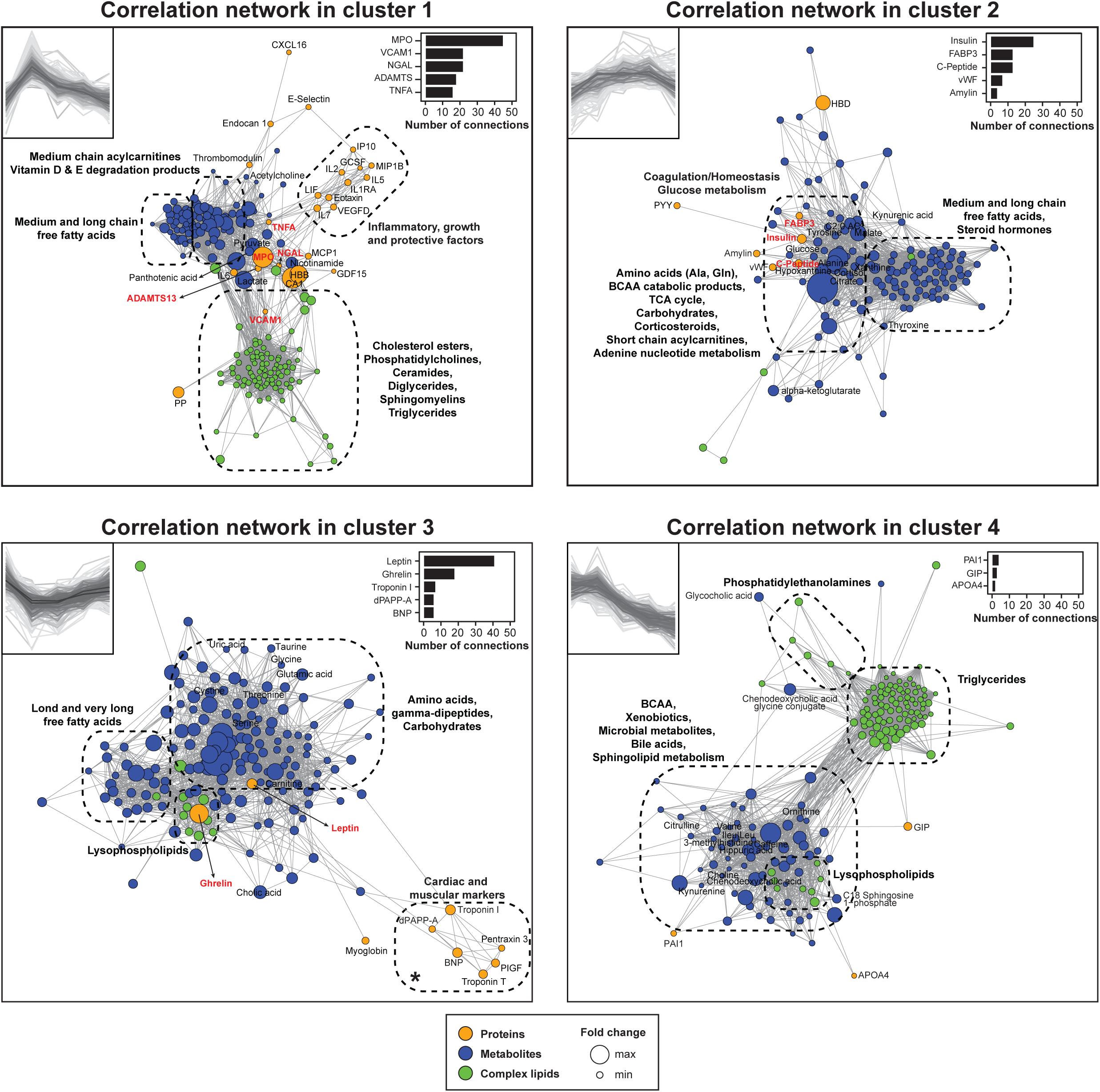
Integrative multi-omic analysis of circulating analytes. Correlation networks of multi-omic measures belonging to each cluster as defined in Figure 2a. Pairwise spearman correlations were calculated in each cluster and were considered significant if Bonferroni was below 0.05. Only the main network was plotted. Nodes were color-coded by molecule type and their size represent the median fold change in response to exercise. The top 5 proteins with the most number of first order connections in each correlation network are displayed. Proteins with more than 10 connections are in bold and red. * Cardiac and muscular markers belong to cluster 3 but the decrease is not significant.

Oxidative stress signaling was apparent through accumulation of myeloperoxidase (MPO) predominantly released from neutrophils via degranulation. Circulating MPO level was strongly associated with neutrophil gelatinase-associated lipocalin (NGAL) (FDR < 1.0E-13), a marker of inflammation (Otto et al., 2015). MPO can further potentiate circulating cytokines by activating endothelial cells to release more cytokines (Odobasic et al., 2016). In the context of exercise, it is believed to signal skeletal muscle damage and recruit macrophages to the damaged sites (Morozov et al., 2006; Reihmane et al., 2012). Interestingly, MPO was among the most responsive molecules (in terms of fold change) and had a central location in the correlation network with the greatest number of connections (n = 45). It was connected to all omic layers bridging inflammatory and growth/protective factors (n = 5) to acylcarnitines (n = 12) and complex lipids (n = 9) suggesting novel roles for MPO in regulating aspects of inflammation and lipid metabolism.

In addition to immune and oxidative stress markers, we detected an increase in molecules that assist in skeletal muscle remodeling such as vascular cell adhesion molecule-1 (VCAM-1), E-selectin and endothelial cell-specific molecule 1 (endocan 1). Similarly to MPO, VCAM-1 was connected to a variety of molecules (n = 22) including neutrophils inflammatory and oxidative stress markers (MPO and NGAL), glycolytic products (lactate and pyruvate), pantothenic acid and many complex lipids (n = 14).

Fatty acid oxidation (FAO) was activated by exercise as indicated by the early accumulation of many acylcarnitines (n = 18) and free fatty acids (n = 30) (**Figure 2d**, **Table S4,8**). Interestingly, our dense sampling revealed distinct trajectories depending on fatty acid composition. Medium chain acylcarnitines (*i.e.* C6:0, C8:0, C10:0, C12:0) accumulated the most following exercise and returned to baseline by 15-30 min in recovery whereas others (*i.e.* C6:1, C8:1) accumulated to a lesser extent and returned to baseline more slowly (30 min to 1 hour) (**Figure 2e**). The increased abundance of circulating medium chain acylcarnitines reflects partial FAO in skeletal muscle (Lehmann et al., 2010; Zhang et al., 2017). The level of circulating free carnitine demonstrated an inverse trajectory (cluster 3) since it is used to form acylcarnitines from free fatty acids. Free fatty acids exhibited three main trajectories with some reaching a maximum at 2 min post-exercise (10-12 carbons, cluster 1), others at 15 min (14-18 carbons, cluster 2) and the remaining decreasing at 2 min in recovery (20-24 carbons, cluster 3) (**Figure 2f**). These observations suggest that long chain fatty acids, in particular free fatty acids with 20-22 carbons (C20:1, C20:2, C22:1, C22:2 and C22:3), are preferentially oxidized during exercise to produce energy resulting in a decrease of these molecules in circulation while partial FAO results in an increased abundance of medium chain fatty acids.

Exercise was also accompanied by a transient accumulation of diverse complex lipids including cholesteryl esters (CE, n = 20), phosphatidylcholines (PC, n = 23), diacylglycerols (DAG, n = 10), ceramides (CER, n = 9) and sphingomyelins (SM, n = 8) (**Figure 2d**). Sphingolipids, and in particular ceramides, may be involved in signaling inflammation in response to exercise as it was observed in the context of inflammatory diseases (Maceyka and Spiegel, 2014).

#### Cluster 2

Molecules in cluster 2 (n = 145) presented a delayed increase post-exercise and a large proportion of these molecules were associated with carbohydrate metabolism. Exercise triggered the secretion of numerous hormones including steroid and thyroid hormones as well as corticosteroids to restore homeostatic balance (**Figure 2d**, **Figure 3b**). Interestingly, correlation networks informed on regulatory mechanisms of hormone secretion in response to exercise. In particular, we detected an increase of cortisol, which in part stimulates the release of hepatic glucose that in turn triggers insulin secretion from pancreatic β-cells to allow cellular absorption (Kjaer, 1998). This was verified in our data with a significant positive correlation between glucose and insulin levels (rho = 0.44, FDR = 1.30E-05). These changes were concomitant with an accumulation of fatty acid binding proteins 3 and 4 (FABP3 and FABP4) that facilitate glucose and free fatty acid uptake in skeletal muscle and heart tissue (Kusudo et al., 2011). Insulin was the most connected proteomic feature in the cluster (n = 25) and was highly correlated with the proinsulin C-peptide and amylin (co-secreted with insulin). Increased glucose metabolism correlated with TCA cycle constituents (malate, citrate, alpha-ketoglutarate) and resulted in a marked increase of products of adenine nucleotide catabolism (*i.e.* hypoxanthine and xanthine) that are markers of ATP turnover (Lewis et al., 2010) (**Figure 2b**). In addition, we detected a delayed increase of the purine end-product uric acid, presumably due to increased synthesis and decreased renal excretion (Sutton et al., 1980) (**Figure S5**). We also detected an increase of coagulation and hemostasis factors (von Willebrand factor (vWF) and A Disintegrin and Metalloprotease with ThromboSpondin motif repeats 13 (ADAMTS-13)) in response to the shear stress induced by treadmill exercise (Stakiw et al., 2008).

#### Cluster 3

Cluster 3 contained molecules (n = 168) that decreased in response to exercise and returned to baseline within 1 hour in recovery. The correlation network was centered around two metabolic hormones leptin and ghrelin suggesting a role in regulation of appetite by exercise (**Figure 3c**). A large proportion of metabolites were amino acids reflecting a central role in exercise physiology (**Figure S4**). Out of 20 proteinogenic amino acids, 15 were significantly changed in response to exercise (FDR < 0.05) (**Figure 2e**). Six amino acids (*i.e.* glutamic acid, cystine, tryptophan, serine, threonine and glycine) belonged to cluster 3 suggesting that they were catabolized by skeletal muscle cells to produce energy (Henriksson, 1991) and re-synthesized in the recovery phase. Four amino acids (*i.e.* alanine, tyrosine, glutamine and proline) presented an opposite trajectory (cluster 2) accumulating as a product of increased cellular metabolism. These changes were expected with alanine and glutamine being released in plasma due to muscle ammonia detoxification (Lewis et al., 2010).

#### Cluster 4

Molecules in cluster 4 (n = 171) were metabolized in response to exercise but did not return to baseline within the 1-hour recovery phase. Some amino acids presented this trajectory and included branched-chain amino acids (BCAA) leucine, isoleucine and valine (**Figure 2g**). BCAA are essential amino acids that cannot be synthesized by the body and are preferentially catabolized by skeletal muscle (Henriksson, 1991) and used to repair damaged skeletal muscle fibers (Negro et al., 2008). BCAA catabolism was evident with a marked increase in branched-chain ketoacids (cluster 2, **Figure S5**). Many other metabolites had the same trajectory with microbial metabolites (**Figure 2h**), xenobiotics (caffeine metabolism) and bile acids.

Cluster 4 also contained many TAG species indicating continuous hydrolysis to generate free fatty acids used for energy production. While most TAG belonged to this cluster, we identified a subset of TAG that increased transiently following exercise (cluster 1). A close examination of their fatty acid composition revealed that TAG in cluster 1 contained fatty acids with more carbons (*P* = 6.01E-07) and unsaturations (*P* = 8.51E-06) than TAG enriched in cluster 4 (**Figure 2i**). TAG in cluster 1 contained fatty acids with signaling properties including arachidonic acid (AA), eicosapentaenoic acid (EPA) and docosahexaenoic acid (DHA) suggesting that the transient burst might play a role in signaling (AA) or compensating for inflammation (EPA, DHA) (Calder, 2013). In contrast, TAG with shorter and saturated fatty acids may be preferentially used for energy (Ranallo and Rhodes, 1998). Altogether, our dense sampling revealed dynamically and functionally distinct subclasses of TAG.

### Dynamic effect of acute exercise on PBMC gene expression

In addition to circulating plasma molecules, we also investigated PBMC gene expression response to exercise both because immune cells play a critical role in muscle damage repair and regeneration and as a system-wide marker of alterations in gene expression (Gjevestad et al., 2015; Philippou et al., 2012; Radom-Aizik et al., 2009; Ulven et al., 2015). Transcripts presented two main trajectories with up- and down-regulated genes reaching a maximum response at 2 min and rapidly returning to baseline within 30-60 min (**Figure 4a**, **Table S4**). Pathway enrichment analysis using all significant transcripts at each time point (FDR < 0.05) revealed expected and novel pathways associated with exercise (**Figure 4b**, **Table S9**).

**Figure 4.**
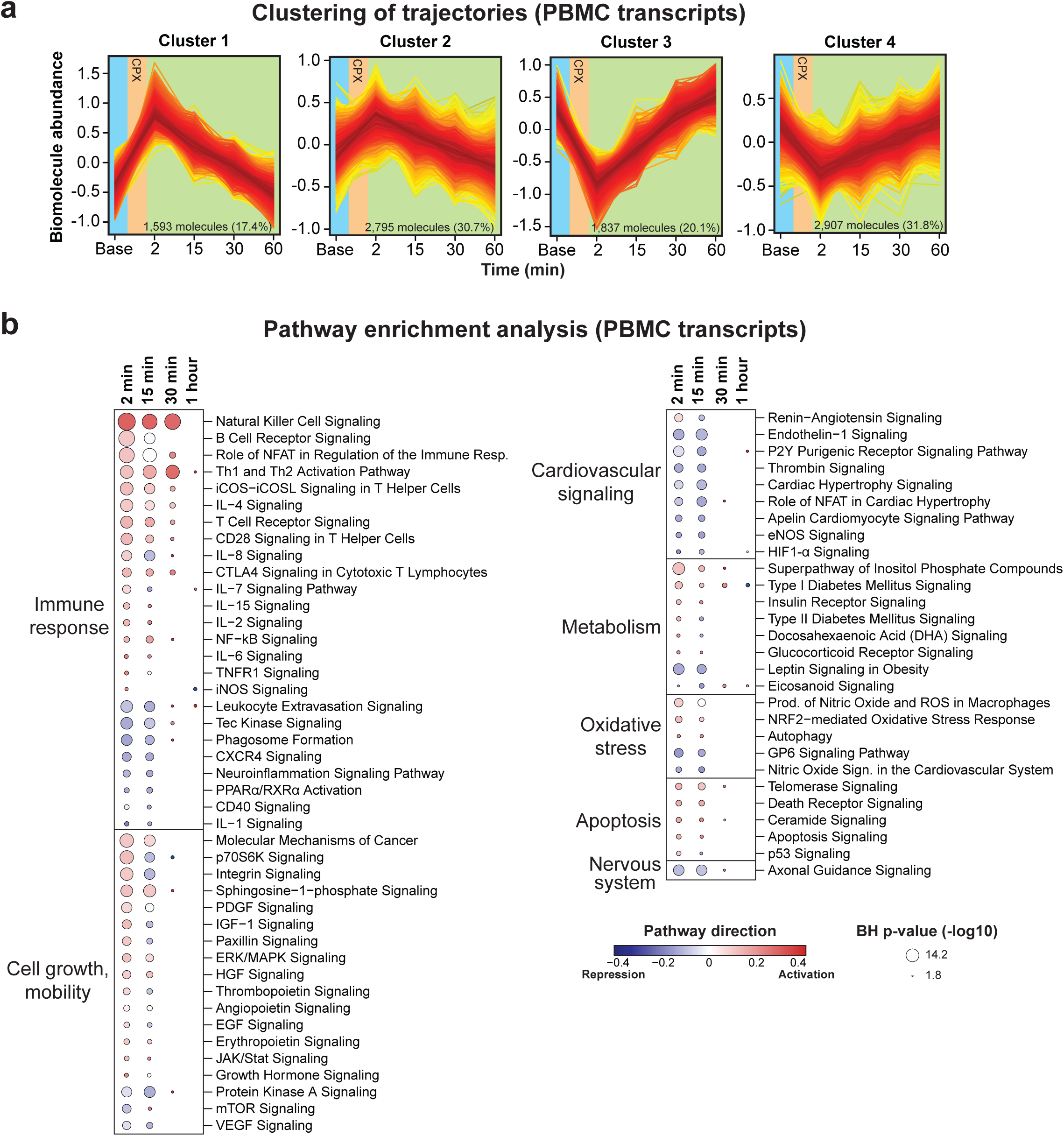
PBMC gene expression changes in response to acute exercise. (**a**) Clustering of longitudinal gene expression trajectories (FDR < 0.05). (**b**) Pathway analysis using PBMC transcripts significantly changing in response to exercise. Significance of pathways was determined by the hypergeometric test (one-sided) in Ingenuity pathway analysis software. *P* values were corrected for multiple hypothesis using the Benjamini-Hochberg method and pathways with FDR below 0.05 were considered significant. Pathway direction is the median log2 fold change relative to baseline of significant transcripts in each pathway (blue: downregulated, red: upregulated). The dot size represents pathway significance.

Exercise induced a robust inflammatory response with an immediate activation of ‘natural killer cell’, ‘Th1 and Th2 activation’, ‘B cell receptor’, ‘T cell receptor’, ‘NF-kB signaling’ and many interleukin signaling pathways as previously reported (Carlson et al., 2011; Connolly et al., 2004; Gjevestad et al., 2015). We also detected activation of pathways related to oxidative stress (*i.e.* ‘production of nitric oxide and reactive oxygen species in macrophages’) and apoptosis (*i.e.* ‘telomerase signaling’, ‘death receptor signaling’). These results are consistent with the early release in circulation of many pro- and anti-inflammatory proteins as well as markers of oxidative stress.

In addition to inflammatory and immune functions, we found a myriad of pathways associated with cell growth and mobility likely involved in muscle tissue repair and remodeling (Connolly et al., 2004). For instance, angiogenesis and wound healing pathways were upregulated in response to exercise (*i.e.* ‘PDGF signaling’, ‘HGF signaling’ and ‘EGF signaling’). Interestingly, we also detected many pathways associated with cardiovascular signaling in PBMCs highlighting the interconnection between exercise and cardiovascular health. These pathways were mainly downregulated and included ‘endothelin-1 signaling’, ‘P2Y purigenic receptor signaling pathway’, ‘thrombin signaling’ and ‘cardiac hypertrophy signaling’. In addition, we also detected dysregulated metabolic pathways including a repression of ‘leptin signaling pathway in obesity’ consistent with a decrease of circulating leptin abundance and activation of ‘superpathway of inositol phosphate compounds’ that may be involved in PBMC activation through synthesis of phosphoinositides (Huang et al., 2007). Although most pathways returned to baseline 30 min post-exercise, some pathways persisted for more than 1 hour (‘Th1 and Th2 activation pathway’) suggesting longer lasting effects of a subset of biological processes. It is known that PBMC number increases with exercise and that the amount of change is cell type-dependent (Millard et al., 2013; Shinkai et al., 1992). Hence, dysregulation of gene expression in response to acute exercise is likely explained by both changes in cell population and in PBMC functional properties.

Finally, we examined allele-specific expression (ASE) in response to exercise and found 11 sites from 9 unique genes with a significant difference in ASE pre- and post-exercise (FDR < 0.05) (**Figure S6**). The strongest associations were found for sites in the human leukocyte antigen (HLA) region (*i.e.* HLA-A, HLA-B, HLA-C, HLA-DPB1, and HLA-DQA1) that were significantly changing following exercise (Booth et al., 2010) (**Table S4**). This suggests that exercise may interact with cis-regulatory variants to modify expression of certain genes.

### Multi-omic analysis of peak VO_2_

Maximum oxygen consumption (peak VO_2_) is among the best predictor of longevity (Ladenvall et al., 2016) and is an important marker of health in the general population and in patients with cardiometabolic disease (Sarullo et al., 2010). In order to gain insights into the molecular underpinnings of healthy molecular profiles, we applied linear regression models to find significant associations with peak VO_2_ at baseline. Associations were also calculated at each time point in recovery to inform on the biological pathways most important to fitness. The models corrected for age, sex, BMI and race/ethnicity that are known factors to impact peak VO_2_ (Kaminsky et al., 2015). The range of peak VO_2_ in the cohort was quite wide with a mean value of 30.6 ± 8.7 ml/kg/min. We found that a large proportion of omic measures associated significantly with exercise capacity at baseline and post-exercise (FDR < 0.05) with 49.6% of complex lipids and 28.0% of metabolites in average across all time points (**Figure 5a**, **Table S10**).

**Figure 5.**
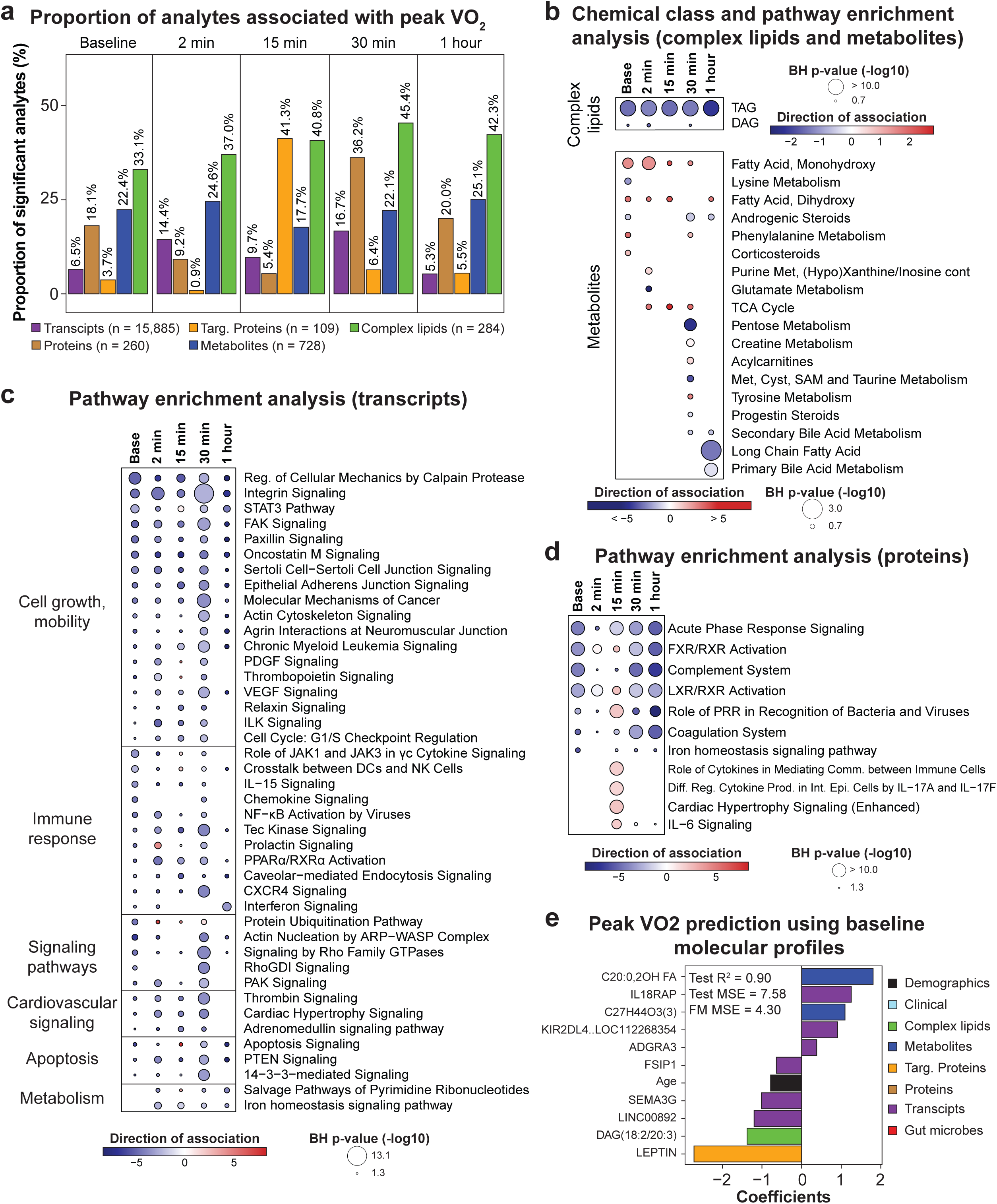
Multi-omic analysis of peak VO_2_. (**a**) Proportion of analytes associated with peak VO_2_ as determined by linear regression analysis after correction for age, gender, body mass index and race/ethnicity. *P* values were corrected for multiple hypothesis using the Benjamini-Hochberg method and analytes with FDR below 0.05 were considered significant. (**b**) Pathway/chemical class enrichment analysis of metabolites and complex lipids using the Kolmogorov–Smirnov method (one-sided). Multiple hypothesis correction was performed using the Benjamini-Hochberg method and pathways with FDR below 0.20 were considered significant. Pathway direction is the median beta coefficient of significant molecules in the pathway (blue: negative association, red: positive association). The dot size represents pathway significance. Pathway analysis using PBMC gene expression (**c**) and circulating proteins (**d**). Significance of pathways was determined by the hypergeometric test (one-sided) in Ingenuity pathway analysis software. *P* values were corrected for multiple hypothesis using the Benjamini-Hochberg method and pathways with FDR below 0.05 were considered significant. Pathway direction is the median beta coefficient of significant molecules in the pathway (blue: negative association, red: positive association). The dot size represents pathway significance. (**e**) Molecules selected in peak VO2 prediction model and associated coefficients. MSE: mean square error.

#### Multi-omic associations at baseline

Leptin correlated the most with exercise capacity at baseline and in recovery (negative association, FDR = 3.1E-06 at baseline) suggesting that total body fat content and/or fat metabolism are key factors of fitness (**Table S10**). Similarly, the transporter of thyroxine and retinol transthyretin (TTR) was positively associated with peak VO_2_ and is a biomarker of lean body mass (Ingenbleek and Bernstein, 2015). Conversely, we detected many TAG and DAG species as well as isoleucine and leucine that negatively associated with exercise capacity (**Figure 5b, Table S11**). Total triglycerides and BCAA are typically associated with poor metabolic health (obesity, prediabetes and type 2 diabetes) (Guasch-Ferre et al., 2016). In contrast, hydroxy-fatty acids and corticosteroids (*i.e.* corticosterone) were positively correlated with peak VO_2_ suggesting that these classes of molecules are markers of health. These findings may be explained by the level of sedentarity of the participants since hydroxy-fatty acids (a marker of FAO) and corticosterone have been shown to increase with exercise training (Droste et al., 2003; Nieman et al., 2013). In addition, we demonstrate that hippuric acid - a marker of gut microbiome diversity as well as fruit and whole grains consumption (Pallister et al., 2017) - was positively associated with fitness. This is consistent with the fact that microbiome diversity is considered an important indicator of overall health (Valdes et al., 2018). We also found some bile acids negatively associated with maximum oxygen uptake (*i.e.* taurocholic acid, tauroursodeoxycholic acid) suggesting either impaired hepatic function (location of conjugation) or gut microbial composition.

PBMC gene expression enrichment analysis revealed numerous pathways negatively associated with fitness that were grouped in the following categories: cell growth and mobility, immune response, signaling pathways, cardiovascular signaling, apoptosis and metabolism (**Figure 5c**, **Table S12**). Even though most pathways were significantly enriched at baseline and at each time point in recovery, many pathways were the most significant 30 min post-exercise suggesting a higher sensitivity at this time point. Importantly, low expression of proteolytic pathways (*i.e.* ‘regulation of cellular mechanics by calpain protease’ and ‘protein ubiquitination pathway’) was associated with better exercise capacity. High activity of the calpain pathway has been shown to play a dominant role in sarcopenia and its decrease correlated with fitness (Bowen et al., 2015). We also found that ‘integrin signaling’ as well as downstream effectors ‘focal adhesion kinase (FAK) signaling’, ‘paxillin signaling’, ‘integrin-linked kinase (ILK) signaling’ and ‘actin cytoskeleton signaling’ were correlated with fitness suggesting an important role of cell adhesion-mediated signaling in exercise. This observation is relevant given the key role of integrin pathway in skeletal muscle health (Graham et al., 2015).

Similarly, many inflammatory pathways were negatively associated with peak VO_2_ such as ‘role of JAK1 and JAK3 in γc cytokine signaling’ ‘IL-15 signaling’ and ‘chemokine signaling’. Metabolic and immune health are tightly interrelated which explains why so many inflammatory pathways are inversely associated with exercise capacity (Hotamisligil, 2006). These findings were validated by circulating proteins with enrichment of inflammatory pathway ‘acute phase response signaling’ as well as complement and coagulation systems (**Figure 5d**, **Table S13**). ‘FXR/RXR activation’ and ‘LXR/RXR activation’ pathways also emerged from our proteomic analysis. Both these pathways regulate glucose and lipid metabolism highlighting their importance in exercise physiology.

#### Multi-omic associations in the recovery phase

Among the molecules and pathways dynamically changed with exercise, a subset associated with peak VO_2_ revealing their biological importance in fitness. We found that skeletal muscle energy plasticity correlated with fitness early in recovery (2-15 min) encompassing positive associations with glucose, malate, lactate and xanthine (**Figure 5b, Table S10,11**). In contrast, tryptophan, cystine, ornithine and allantoin were negatively correlated with peak VO_2_ suggesting a preferential usage of these molecules to fuel the TCA cycle and generate energy. Later in recovery (30-60 min), other biological processes emerged with energy homeostasis represented by a negative association of peak VO_2_ with long chain free fatty acids and medium chain acylcarnitines as well as a positive association with cortisol.

Importantly, our proteomic data revealed a strong link between inflammatory response to exercise and peak VO_2_ in recovery as demonstrated by a significant enrichment of the following pathways ‘role of cytokines in mediating communication between immune cells’ and ‘IL-6 signaling’ (**Figure 5d**). Among pro-inflammatory factors, many interleukins (13 out of 21), members of the TNF superfamily (TNF-β, FASL, CD40L) and interferons (IFN-α and IFN-γ) positively associated with peak VO_2_ 15 min post-exercise (**Table S13**). Intriguingly, the levels of TNF-α and IL-6 were not associated with exercise capacity. Similarly, many protective and growth factors positively (n = 9) and negatively (*i.e.* GM-CSF and GDF15) associated with VO_2_ max. Altogether, these data indicate that a higher level of inflammatory and growth/protective factors at 15 min in recovery is a critical biological process associated with fitness. We also detected proteins negatively associated with peak VO_2_ including regulators of glucose metabolism adipsin and gastric inhibitory polypeptide (GIP) as well as cardiac FABP4.

### Multi-omic prediction of CPX parameters

Next, we evaluated how well baseline multi-omic measurements could predict peak VO_2_, the minute ventilation carbon dioxide production relationship (VE/VCO_2_ slope) and the respiratory exchange ratio (RER). VE/VCO_2_ slope has been shown to be prognostic of heart failure (Arena et al., 2004) and RER is a marker of maximum effort (Albouaini et al., 2007). In addition to the multi-omic data generated in this study, we included clinical laboratory (**Table S14**) and gut microbiome data (**Table S15**) generated within 4 months of the exercise date (54.2 and 112.9 days in average, respectively). We and others have shown that these measurements are fairly stable within this time range (Zhou et al., 2019). First, we identified highly predictive molecules using a Bayesian network algorithm and then we used ridge regression modeling. We built predictive models for peak VO_2_, VE/VCO_2_ and RER with cross-validated R^2^ of 0.90, 0.75 and 0.81, respectively (**Figure 5e**, **Tables S16-18**).

In order to understand the relative importance of each molecule type to the output CPX parameters, we compared predictive models generated from multi-omic data *versus* data from a single ome. We found that peak VO_2_ could be predicted accurately (R^2^ > 0.80) using metabolomic or transcriptomic data only suggesting that metabolites and PBMCs transcripts contain information on the molecular processes that translate into exercise capacity (**Table S16**). Age and BMI were found to be consistently chosen across multiple predictive models demonstrating their importance in fitness level as previously reported (Ribisl et al., 2007). Leptin was confirmed as a critical marker of exercise capacity and other measures such as interleukin 18 receptor accessory protein (IL18RAP) emerged from this analysis.

The model generated to predict VE/VCO_2_ from transcriptomic data alone was superior than any other single ome models (R^2^ = 0.87; **Table S17**). The proportion of Verrucomicrobia phylum in the gut was selected in the multi-omic prediction model (positive association). Verrucomicrobia is generally associated with improved metabolic health and regulation of glucose homeostasis (Dao et al., 2016). RER could be predicted accurately with lipidomic and transcriptomic data only (R^2^ > 0.65) (**Table S18**). As expected, glucose level was important to predict RER and this analysis revealed emerging markers including the proportion of Butyricimonas genus in the gut and plasma eicosapentaenoic acid.

Altogether, we demonstrate the possibility of predicting important physiological parameters using resting molecular measurements, but also suggest potential key molecules and treatments (*e.g.* pre- and probiotics) that may be useful for improving exercise capacity and healthy physiology.

### Differential response to exercise in insulin resistant participants

Participants presented a wide range of peripheral insulin resistance with fourteen participants categorized insulin sensitive (IS) and 16 insulin resistant (IR, SSPG ≥ 150 mg/dl). We investigated the differential response to exercise in IR relative to IS participants with the goal of highlighting abnormal biological processes associated with insulin resistance. Participants from both groups reached a similar RER at peak exercise (*P* = 0.59), indicating similar levels of exhaustion (**Figure 6a**). Peak VO_2_ and VE/VCO_2_ were not significantly different between the two groups despite a trend towards lower exercise capacity in IR participants (*P* = 0.11).

**Figure 6.**
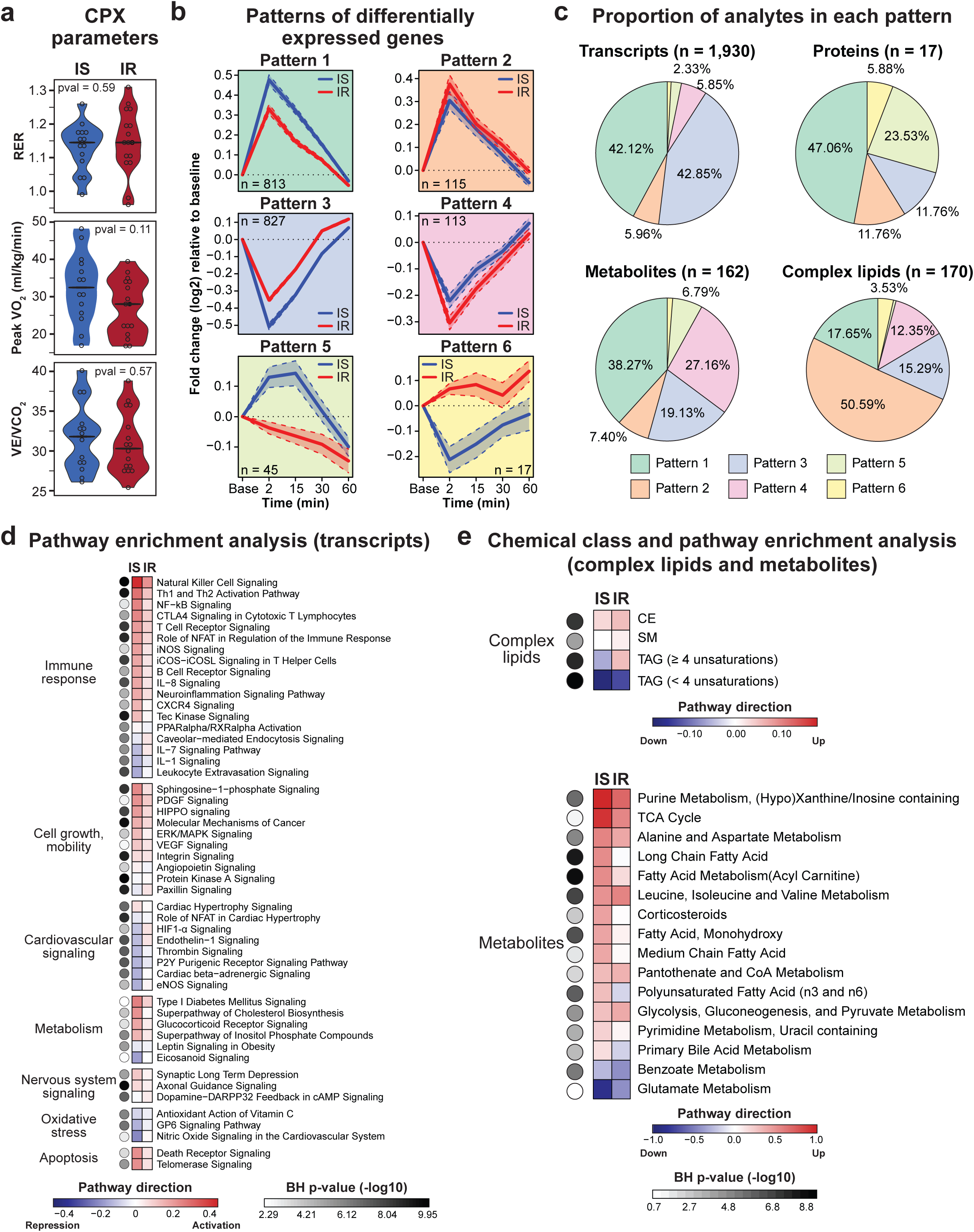
Differential response to acute exercise in insulin resistant participants. (**a**) Violin plots showing CPX parameters in insulin sensitive (IS) and resistant (IR) participants as defined by the modified insulin suppression test (IR: steady-state plasma glucose (SSPG) ≥ 150 mg.dl^-1^). A two-sided Student t-test (normal distribution) or a Wilcoxon rank sum test was used for differential analysis. The violin plots illustrate the kernel probability density and the horizontal bar depicts the median of the distribution. (**b**) Patterns of differentially expressed genes in IS and IR participants. A linear mixed model correcting for personal baseline, age, sex, body mass index and race/ethnicity was employed. *P* values were corrected for multiple hypothesis using the Benjamini-Hochberg method and transcripts with FDR below 0.05 were considered significant. The solid line represents the mean and the dashed line represents the 95% confidence interval. (**c**) Pie charts depicting the proportion of significant transcripts (FDR < 0.05), proteins (FDR < 0.20), metabolites (FDR < 0.10) and complex lipids (FDR < 0.20) in each of the six patterns as defined in (**b**). (**d**) Pathway analysis using PBMC gene expression. Significance of pathways was determined by the hypergeometric test (one-sided) in Ingenuity pathway analysis software. Multiple hypothesis correction was performed using the Benjamini-Hochberg method and pathways with FDR below 0.05 were considered significant. Pathway direction is the median of max/min fold change relative to baseline of significant molecules in the pathway (blue: downregulated, red: upregulated). The color of the dots represents pathway significance. (**e**) Pathway/chemical class enrichment analysis of metabolites and complex lipids using the Kolmogorov–Smirnov method (one-sided). *P* values were corrected for multiple hypothesis using the Benjamini-Hochberg method and pathways with FDR below 0.20 were considered significant. Pathway direction is the median of max/min fold change relative to baseline of significant molecules in the pathway (blue: downregulated, red: upregulated). The color of the dots represents pathway significance.

Linear mixed models were used and corrected for personal baseline, age, sex, race/ethnicity and BMI. We found 2,279 differential analytes across all omic datasets and categorized them based on their longitudinal trajectories into 6 patterns (**Table S19**). Patterns generated from PBMC transcripts are presented in **Figure 6b**, whereas results from proteins, metabolites and complex lipids are presented in **Figure S7**. Patterns 1 and 2 contained upregulated analytes with a higher amplitude in IS and IR participants, respectively. Analytes in patterns 3 and 4 were downregulated and patterns 5 and 6 presented opposite trajectories. As an example, 1,930 PBMC genes were differentially expressed in IR participants (FDR < 0.05) with most genes belonging to patterns 1 and 3 (84.97% in total) (**Figure 5c**) suggesting a dampened response of immune cells.

This observation was verified at the pathway level with generally a stronger response in IS subjects (**Figure 6d**, **Table S20**). Differential pathways belonged to various biological functions including inflammatory, cell growth and mobility, cardiovascular and metabolism. Differential inflammatory response in IR participants was evident with a milder activation of ‘natural killer cell signaling’ and ‘Th1 and Th2 activation pathway’. Similarly, activation of apoptosis (‘telomerase signaling’) and cell growth and mobility pathways (‘PDGF signaling’) were stronger in IS. These findings are consistent with other studies describing a diminished PBMC response to weight gain (Piening et al., 2018), viral infection and immunization (Zhou et al., 2019) in IR subjects. Circulating proteins reinforced gene expression results with a smaller amount of MPO and NGAL immediately after exercise in IR individuals (**Figure S7c**). Interestingly, the inflammatory protein TNF-α had a similar amplitude of response in both groups but persisted for a longer time in IR and ultimately returned to baseline levels 1 hour post-exercise in comparison to 15 min for IS subjects (**Figure S7d**). The same pattern was observed for IL-6 suggesting abnormal cytokine re-absorption or neutralization in IR participants. More dramatically, pentraxin 3, a marker of acute inflammatory response, increased immediately post-exercise in IS whereas its level remained low in IR across the whole study (**Figure S7c**). We observed higher levels of this inflammatory factor at baseline in IR participants which may explain this difference. Interestingly, some cardiovascular pathways involved in vasoconstriction (*i.e.* ‘endothelin-1 signaling’), response to hypoxia (*i.e.* ‘HIF-1α signaling’) and nitric oxide synthesis (*i.e.* ‘eNOS signaling’) were impaired in IR with a mild upregulation relative to a repression in IS. We also observed an opposite response for troponin T with an immediate release in circulation in IS and a delayed increase in IR reaching similar levels 1 hour post-exercise.

Although most genes responded in the same direction, some responded in opposite directions (pattern 5 and 6). Pathway enrichment analysis revealed that ‘protein ubiquitination pathway’ (FDR = 3.9E-06) was upregulated in IS and downregulated in IR. Protein ubiquitination is a key mechanism involved in protein turnover regulation in skeletal muscle following exercise (Cunha et al., 2012). These results suggest that IR individuals have altered proteasome activity in PBMCs.

Similar analysis using circulating metabolites revealed that many major biological pathways impacted by exercise were abnormal in IR participants (**Figure 6e**, **Table S21**). For instance, lipid (increase of acylcarnitines and medium chain free fatty acids), carbohydrate (increase of glucose) and amino acid metabolism (decrease of those used for energy production) responded more strongly in IS resulting in the accumulation of malate (TCA cycle), lactate, hypoxanthine and xanthine (ATP turnover) and alanine (muscle ammonia detoxification) (**Figure S7e**). Consistent with these observations, metabolic hormones including gastric inhibitory polypeptide (GIP), leptin and ghrelin decreased more strongly in IS following exercise (**Figure S7c**). In contrast, insulin was among the few molecules that responded more strongly in IR participants. In addition, insulin secretion was delayed reaching a maximum 15 min post-exercise in IR versus 2 min in IS. This was expected since a higher amount of insulin is necessary for peripheral tissues to absorb glucose in circulation. Interestingly, we observed that the cortisol response had a similar amplitude in IS and IR but returned to baseline 1 hour post-exercise in IR whereas it remained high in IS individuals. In addition, we observed an impaired energy homeostasis with insulin resistance. Long and polyunsaturated free fatty acids are oxidized during exercise to produce energy and are resynthesized in recovery. The level of these molecules increased by 15 min post-exercise and remained high until 30 min in IS subjects whereas their level stayed low in IR participants suggesting an abnormal synthesis of free fatty acids.

In contrast with transcripts, proteins and metabolites, most complex lipids (50.59%) accumulated more strongly in IR individuals following exercise (pattern 2) (**Figure 6c**). In particular, CE and SM tended to accumulate more immediately post-exercise in IR group (**Figure 6e**, **Table S21**). Interestingly, the increase of unsaturated TAG (4-12 unsaturations) post-exercise was only visible in IR participants but not in IS. On the other hand, saturated TAG (0-3 unsaturations) decreased more in IS suggesting a more efficient hydrolysis.

### Multi-omic outlier analysis to highlight personal abnormalities

In order to gain molecular insights at the individual level, we examined the number of outlier molecules (FDR < 0.05) at baseline (absolute levels) and in response to exercise (delta) across multi-omic datasets in each individual in the cohort. Despite all the participants were considered healthy, four individuals presented outlier molecular profiles (**Figure S8a**). These participants were not outlier for any demographic, echocardiography, vascular ultrasound, CPX or clinical parameters (**Figure S8b**). Individuals ZLTUJTN and ZJBOZ2X had significantly more transcript outliers than the rest of the cohort at baseline. Interestingly, these participants did not display many outliers in the other omes suggesting that these results were not due to sample quality/integrity. Pathway enrichment analysis revealed that ‘interferon signaling’ (FDR = 3.2E-10), ‘activation of IRF by cytosolic pattern recognition receptors’ (FDR = 3.9E-09) and ‘protein ubiquitination pathway’ (FDR = 2.7E-05) were abnormal in individual ZLTUJTN (**Table S22**). In addition, ‘iron homeostasis’ (FDR = 1.4E-03) and ‘heme biosynthesis’ (FDR = 4.9E-03) were abnormal in this individual who also was an outlier for the clinical labs with low mean corpuscular volume (MCV, 72.8 fl per cell) and mean corpuscular hemoglobin (MCH, 22.6 pg/cell) as well as high red cell distribution width (RDW, 20.5%) (FDR < 0.05). This participant was also mildly anemic (hemoglobin 13.4 g/dl) and a subsequent clinical work-up of these findings revealed that he was an alpha thalassemia carrier. Interestingly, the same participant also presented many transcript outliers in response to exercise with ‘hypoxia signaling in cardiovascular system’ (FDR = 3.8E-05) and ‘iNOS signaling’ (FDR = 2.3E-04) which may also relate to his microcytic anemia/alpha thalassemia carrier status (Ho et al., 2012). Participant ZL63I8R presented an outlier protein profile at baseline and in response to exercise consisting of immune proteins, growth factors and proteins involved in hemostasis, blood clot and lipid metabolism (**Figure S8c**). These results show that individuals can present large differences in their molecular composition and that deep individual molecular profiles may be useful and sensitive to detect subclinical conditions.

## Discussion

Fitness represents one of the strongest predictors of survival in the general population and in patients with established cardiovascular disease (Blair et al., 1989; Myers et al., 2002). The major contribution of our study is to provide an integrated system-wide molecular choreography following an acute bout of exercise across the spectrum of insulin sensitivity and resistance. The dense sample collection pre- and post-exercise allowed a very fine mapping of co-occurring changes highlighting the complex interplay between systems. We build on the background of several clinical and experimental exercise studies by integrating dynamic multi-omic changes measured in immune cells (PBMCs) and plasma in response to a symptom-limited cardiopulmonary exercise (Lewis et al., 2010; Price et al., 2017).

The molecular response to exercise was not only well orchestrated but also showed a high degree of individuality. Our study reveals a plethora of biological processes involved in exercise spanning multiple key compartments (*i.e.* skeletal muscle, adipose tissue, immune cells, liver and cardiovascular system) as well as their complex interplay during and after exercise. The main biological functions captured relate to energy metabolism, inflammation, oxidative stress and homeostasis as well as tissue repair and remodeling. We detected expected changes in plasma including an accumulation of metabolites related to glucose oxidation (*i.e.* malate, pyruvate and lactate) and pro-inflammatory factors (IL-6 and TNF-α). Previous studies had reported a higher fold change in lactate as these studies focused on higher intensity and more sustained exercise often performed in younger individuals (Allen et al., 1985; Oyono-Enguelle et al., 1990). We also demonstrate that several hormones (*i.e.* cortisol, insulin, leptin and ghrelin) likely play a key role in the regulation of exercise recovery. These hormones are involved in hepatic glucose release, cellular glucose absorption as well as regulation of appetite. Interestingly, we also observed an increase of neuroactive metabolites acetylcholine and kynurenic acid (KYNA) in response to exercise which links physical activity to mental health. KYNA is a product of tryptophan metabolism, is produced in exercise-stimulated skeletal muscle and has been shown to deliver antidepressant activity (Agudelo et al., 2014).

Our study also adds granularity on the potential role of subset species of acylcarnitines, free fatty acids, complex lipids and amino acids. Depending on their number of carbons and unsaturations, free fatty acids were divided among different time clusters with longer chain fatty acids decreasing as they oxidize to provide energy. In contrast, medium chain free fatty acid and by extension medium chain acylcarnitines accumulated reflecting partial FAO in skeletal muscle. We have also found that the dynamic of some complex lipids was dependent on their carbon and unsaturation levels. Even though most triglycerides (TAG) decrease in recovery presumably used for energy production (Ranallo and Rhodes, 1998), a subset containing long-chain poly-unsaturated fatty acids increased immediately post-exercise (cluster 1). This finding may reflect pro- or anti-inflammatory signaling since these TAG contain arachidonic acid, eicosapentaenoic acid and docosahexaenoic acid (Calder, 2013). We also detected other complex lipids in cluster 1 including cholesterol esters, phosphatidylcholines, ceramides, diglycerides and sphingomyelins. As part of the orchestrated lipid response, we observed an increase in free fatty acid binding proteins 3 and 4 (FABP3 and FABP4) which facilitate glucose and free fatty acid uptake in skeletal muscle and heart tissue (Kusudo et al., 2011). FABP3 has been proposed as an early marker of myocardial ischemia (Tanaka et al., 1991) and the observation of its increase post-exercise invites further investigation on its potential role during exercise testing. Amino acids also presented different temporal profiles depending on their nature differentiating essential and non-essential amino acids as well as those catabolized for energy generation and those produced due to heightened cellular metabolism.

One of the strengths of our study was to highlight the complex crosstalk between oxidative stress, inflammation and metabolism during exercise recovery. Myeloperoxidase (MPO) - a marker of oxidative stress secreted by neutrophils - was one of the proteins that increased the most in response to exercise and had a central location in the acute response network bridging inflammatory and metabolic factors. For example, the circulating level of MPO was strongly associated with neutrophil gelatinase-associated lipocalin (NGAL) which marks inflammation. NGAL is an emerging marker of acute renal injury in a different context and its complementary role to MPO needs to be further explored (Goodsaid et al., 2009). To a lesser extent, growth differentiating factor-15 (GDF-15), which is a strong marker of survival in cardiometabolic disease (Wallentin et al., 2013), was also increased in the early phase of exercise recovery. Although transcriptomic analysis was performed on peripheral blood mononuclear cells (PBMCs), we could detect changes far beyond the pathways of immunity and inflammation including tissue repair and remodeling, cardiovascular health such as nitric oxide and endothelin-1 signaling, vascular and epithelial growth factors and apoptosis to highlight a few. Tissue repair and remodeling was evident via the upregulation of IGF-1 signaling pathway that activates by phosphorylation serine/threonine kinases (p70S6K) necessary to increase protein synthesis and rebuild the damaged muscle (Schiaffino and Mammucari, 2011).

Multi-omic associations with exercise capacity bring novelty on the molecular signature of fitness. Prior to our study, major insights into the integrated omic of exercise were based on small experimental animal studies. For example, important biological insights were obtained in rats from low (LCR) and high capacity runners (HCR) (Koch and Britton, 2001). The HCR rats had lower visceral adiposity, lower fasting glucose and insulin as well as increased natural life span (Koch et al., 2011). The landmark study of Overmyer *et al*. (Overmyer et al., 2015) demonstrated that HCR efficiently oxidize fatty acids and branched-chain amino acids sparing glycogen and reducing accumulation of short- and medium-chain acylcarnitines. HCR rats also demonstrated enhanced mitochondrial protein deacetylation with exercise, which appeared to augment this efficient fuel use. Ren et al. (Ren et al., 2016) further showed that several pathways associated with mitochondrial function, cell adhesion and extracellular matrix were enriched in skeletal muscles that differ between HCR and LCR lines.

In our study of humans, we found that skeletal muscle energy plasticity was a critical aspect of fitness with an efficient i) energy metabolism evident via the extent of accumulation of malate, lactate and xanthine early in recovery (2-15 min) and ii) energy homeostasis shown by the rate of re-synthesis of long chain fatty acids later in recovery (30-60 min). We also found that participants metabolic health was determinant in exercise capacity including baseline abundance of leptin and triglycerides. Besides metabolism, our broad multi-level profiling identified inflammation and vascular growth and signaling as important modulators of exercise capacity. At the gene expression and circulating protein levels, baseline chronic inflammation was negatively associated with peak VO_2_ whereas pro-inflammatory response 15 min in recovery was found to be key to fitness. In addition, particular attention needs to be given to the two most significant pathways from PBMC transcripts, *i.e.* calpain and integrin pathways. Calpain pathway is involved in muscle regulation and has been associated with sarcopenia, an important mechanism of frailty in our aging population. The integrin pathway mediates the control of insulin-like growth factor receptor (IGF1R) signaling and in turn regulates the muscular response to exercise (Legate et al., 2009). For example, α7β1 integrin subunit was found to be protective of muscle damage in response to strain associated with eccentric lengthening in mice (Mahmassani et al., 2017). This subunit is also known to be expressed on cardiac myocytes (Carrier et al., 1991). We further demonstrated that complex physiological status such as fitness could be predicted from 10-12 multi-omic measures at rest, suggesting that exercise testing parameters could be calculated from resting blood-based analytes in the future. Interestingly, some microbial strains were selected in these models suggesting important roles of the gut microbiome as it was recently demonstrated with the performance-enhancing microbe genus Veillonella (Scheiman et al., 2019).

Our study also demonstrates that exercise profiling can provide insights into the pathophysiology of metabolic conditions. Given the epidemic of obesity, understanding deleterious effects of insulin resistance becomes critical (Kahn et al., 2006). The combination of dense longitudinal sampling and statistical analysis revealed key differences between insulin sensitive and resistant individuals which would not be apparent at baseline. Despite the small sample size, significant differences were noted in all the main biological processes impacted by exercise including inflammatory, oxidative stress, vascular, hypertrophic and cell growth pathways. In addition to a reduced inflammatory response in insulin resistant participants (often related to a higher baseline activation), we observed a reduced efficiency to oxidize free fatty acids, produce energy and restore energy homeostasis. We also detected differential response in glutamate metabolism which is implicated in coronary heart disease (Qi et al., 2013). Furthermore, cardiovascular signaling showed marked differences between groups of insulin sensitivity and included, endothelin-1, a vasoconstrictor and therapeutic target in vascular biology (Marasciulo et al., 2006), thrombin, a critical enzyme in the coagulation pathway (Yanagisawa, 1994) and cardiac beta-adrenergic signaling found to be reduced in heart failure syndrome or autonomic disorders (Lymperopoulos et al., 2013). Omic profiling across the spectrum of insulin resistance demonstrates the potential of using exercise to identify mediators or biomarkers of disease. For example, the study of Lewis et al. used exercise profiling to identify novel metabolites (indoleamine 2,3-dioxygenase (IDO)-dependent tryptophan metabolites) associated with right heart function during exercise testing (Lewis et al., 2016).

To move the field of precision exercise omics, one has to better understand the normal reference range. Our outlier analysis represents one example where pathways dysregulation can help identify potential abnormal processes. Since the exercise response shows high individuality - personalized exercise profiling could prove to be valuable and more sensitive for disease detection with the logistic challenges of creating an easy clinical pipeline.

This work should be assessed in the context of its limitations. Our cohort was small (n = 36) and caution should be given to the generalization of the results due to the self-selection of participants and generally large heterogeneity in humans. However, to our knowledge our deep phenotyping combined with a personalized exercise test is one of the most comprehensive molecular studies ever performed. Importantly, most of the data are open access. The interactions described are in the context of a single bout of acute exercise, and we encourage future studies to assess the augmentation of our described responses with different forms of training. While our study advised the limitation of activity prior to testing, by design we demonstrated that physical activity performed in the community, may have a greater acute effect than the underlying disease state being investigated. As further attempts are made in integrating an array of longitudinal clinical and molecular signals, vigilance will be required in standardization in performance testing and within the data of the individual. Other large initiatives underway, of note, Molecular Transducers of Physical Activity Consortium (NCT03960827) will scale the concepts presented here, among a diverse group of participants.

In conclusion, our study provides an in-depth and integrated multi-omic profiling of exercise response. The translation potential of the study resides in the discovery of promising resting biomarker signature of fitness as well as in demonstrating the value of exercise molecular testing in identifying key differences in the mechanisms of insulin resistance. Ongoing studies will help refine reference range in exercise omic response that will guide future population and personalized outlier analysis.

## Acknowledgements

This work was supported by grants from the National Institute of Health (NIH) Human Microbiome Project (HMP) 1U54DE02378901, S10OD020141 as well as SDRC grant P30DK116074. K.M. received support from an Australian Government Research Training Program (RTP) Scholarship. We thank Yael Rosenberg-Hasson and the Human Immune Monitoring Center (HIMC) at Stanford for generating targeted proteomic data. We also would like to thank the iPOP participants who generously gave their time and biological samples.

## Author Contributions

K.C., M.P.S., F.H., K.M., J.W.C and S.M.S-F.R contributed to the conceptualization. K.C., S.W, F.H., M.P.S., K.M., J.W.C. and D.P. contributed to the methodology. K.C. (targeted proteomics), K.C., B.L.M., S.C. and M.E. (metabolomics), D.H., M.E. and K.C. (lipidomics), S.A., J.V.Q. (proteomics), M-S.T., E.W., B.E., H.C. (transcriptomics) contributed to omics generation and/or processing. K.M., F.H. and J.W.C contributed to cardiovascular and CPX data collection. K.C., S.W., K.M., J.W.C., S.M.S-F.R., M.A., Y.K. and W.Z. contributed to data curation. K.C., S.W., F.H., K.M., D.H. and M.P.S. contributed to data visualization. K.C., S.W., K.M., D.H., A.A.M., E.W., S.A., B.B, M.D. and D.A.K contributed to formal analysis. A.B. and K.C. contributed to data deposition. K.C., F.H. and M.P.S. contributed to project administration. M.P.S. and F.H. contributed to supervision. K.C., F.H., K.M., M.P.S., J.W.C. and S.T. contributed to writing and preparing the original draft.

## Data availability

Raw and processed omic data (transcriptome, targeted proteome, proteome, metabolome, lipidome) are hosted on the NIH Human Microbiome 2 project site (https://portal.hmpdacc.org/) under the T2D project. Microbiome and clinical laboratory data have been provided in the Supplementary Data files.

## Material and methods

### Participant Recruitment and IRB Consent

Study participants were enrolled as “healthy volunteers” in the framework of the NIH integrated Human Microbiome Project 2 (iHMP) (Zhou et al., 2019). Inclusion and exclusion criteria are described in detail elsewhere (Schussler-Fiorenza Rose et al., 2019). Among the iHMP cohort, 36 subjects provided informed written consent to participate in the exercise study under a research study protocol approved by the Stanford University Institutional Review Board (IRB 23602). Participants were screened for contraindications for exercise testing and comorbidities with a basic health questionnaire. Participant demographics can be found in **Table 1**. Thirty out of 36 participants underwent the modified insulin suppression test to determine steady-state plasma glucose (SSPG) levels as described (Schussler-Fiorenza Rose et al., 2019) and classify the participants as insulin sensitive (n = 14, SSPG < 150 mg/dl) and insulin resistant (n = 16, SSPG ≥ 150 mg/dl). The remaining six individuals didn’t perform the test because of medical contraindications.The cohort was composed of normoglycemic (n = 16), prediabetic (n = 16) and diabetic (n = 4) individuals as determined by fasting plasma glucose (FPG) and hemoglobin A1C (HbA1C) levels measured within 2 months of the exercise date (prediabetic range: 100 mg/dl ≤ FPG < 126 mg/dl or 5.7% ≤ HbA1C < 6.5%; diabetic range FPG ≥ 126 mg/dl or HbA1C ≥ 6.5%).

### Study Design

Overnight-fasted participants (10-12 hours) arrived at Stanford Clinical Translational Research Unit (CTRU) at 7:00 am in the morning. Resting vital signs including heart rate, blood pressure, oxygen saturation, height and weight as well as blood glucose were recorded. Blood was collected from participants at baseline (7:15 am) and echocardiography as well as vascular ultrasound were performed at rest (7:45 am). Afterwards, the study subjects underwent symptom-limited cardiopulmonary exercise (CPX) testing (8:00 am) and received a stress echocardiography. Additional blood samples were collected longitudinally post-exercise.

### Transthoracic Echocardiography

Participants underwent transthoracic echocardiography using commercially available echocardiographic systems (iE33; Philips Medical Imaging, Eindhoven, the Netherlands). Post-stress images were acquired immediately post-exercise, as per international consensus guidelines and all participants had satisfactory imaging. Digitized echocardiographic studies were analyzed on Xcelera workstations in accordance with published guidelines of the American Society of Echocardiography (ASE) (Lang et al., 2015). Left ventricular diameters were indexed on height, while mass and volumes were indexed on body surface area. Left ventricular ejection fraction (LVEF) was calculated by modified biplane Simpson’s rule of apical imaging (Wilson et al., 1998). Left ventricular global longitudinal strain (LV GLS) was calculated from apical imaging on manual tracings of the mid wall with the formula for Lagrangian Strain % = 100 x (Lt - L0)/L0), as previously described (Smith, 2016). With tissue doppler imaging (TDI), we used peak myocardial early diastolic velocity at the lateral mitral annulus and the assessment of trans mitral to TDI early diastolic velocity ratio (E/e’) (Lee et al., 2010; McClelland et al., 2015). Left atrial volume was calculated by the biplane disk summation technique and indexed to body surface area as described by the ASE (Lang et al., 2015).

### Vascular Ultrasound

Screening for subclinical atherosclerosis was performed using vascular ultrasound of the carotid and femoral arteries with a 9.0 MHz Philips linear array probe and iE33 xMATRIX echocardiography system (Philips, Andover, MA, USA). Vascular stiffness was assessed using central pulse wave velocity (PWV). PWV was calculated with the formula distance (m)/transit time (s) by assessing the flow of the carotid and femoral arteries separately and normalizing with electrocardiogram (Calabia et al., 2011).

### Symptom-limited Cardiopulmonary Exercise Testing

Symptom-limited cardiopulmonary exercise (CPX) testing was performed with an individualized ramp treadmill protocol (Myers et al., 1991). Participants were encouraged to exercise to maximal exercise capacity with a target duration of 8-12 minutes following the ramp protocol tuned to individuals exercise capacity as determined by a questionnaire. All participants ceased exercise due to dyspnea and/or fatigue and none experienced chest pain or terminated the study due to arrhythmia. Ventilatory efficiency (VE), oxygen consumption (VO_2_), volume of carbon dioxide production (VCO_2_) and other CPX variables were acquired breath-by-breath and averaged over 10 second intervals (Omnia CPET, CosMed USA, Concord, CA, USA). A respiratory exchange ratio (RER; VCO_2_/VO_2_) > 1.05, heart rate (HR) > 85% of predicted maximum and Rating of Perceived Exertion (RPE6-20; Borg Perception, Hasselby, Sweden) were determined to indicate peak effort. Peak oxygen uptake VO_2_ was calculated as the highest VO_2_ levels. VE and VCO_2_ responses throughout exercise were used to calculate the VE/VCO2 slope via least squares linear regression (y = mx + b, m = slope) (Arena et al., 2003).

### Blood Collection and Sample Preparation

Intravenous blood from the upper forearm was drawn from overnight-fasted participants at baseline (before exercise) as well as 2 min, 15 min, 30 min, and 1h post-exercise. Samples at baseline and 1-hour time points were collected in the CTRU while samples collected 2 min, 15 min and 30 min post-exercise were collected in the exercise laboratory. For time points 2 min, 15 min and 30 min, the intravenous cannula was flushed with 5 ml of normal saline after each blood draw to prevent obstruction and contamination. Specimens were immediately placed on ice after collection to avoid sample deterioration and processed together immediately after collection of the last sample (9:00 am). Blood was collected in a purple top tube vacutainer (BD), layered onto Ficoll media and spun at 2,000 rpm for 25 min at 24°C. The top layer EDTA-plasma was pipetted off, aliquoted and immediately frozen at −80°C. The peripheral blood mononuclear cells (PBMC) layer was collected, counted via cell counter and aliquots of PBMCs were further pelleted and flash-frozen. Multi-level molecular profiling was performed on all blood samples including gene expression from PBMCs (transcriptomics), proteins (targeted and untargeted proteomics), metabolites (untargeted metabolomics), and complex lipids (semi-targeted lipidomics) from plasma. Transcriptomics, metabolomics and targeted proteomics were performed on fresh EDTA-plasma aliquots while untargeted proteomics and lipidomics were performed on EDTA-plasma that went through one freeze-thaw cycle.

### RNA-sequencing from peripheral blood mononuclear cells (PBMCs)

#### RNA extraction and library preparation

The transcriptome was evaluated by RNA sequencing (RNA-seq) from bulk PBMCs. PBMCs were thawed on ice, and subsequently lysed and processed to DNA, RNA and protein fractions using silica-membrane spin columns from the AllPrep DNA/RNA/Protein kit (cat# 80004, Qiagen, Chatsworth, CA, USA). PBMCs were processed in a randomized order. A Bravo NGS Workstation (Agilent, Santa Clara, CA, USA) was used to perform automated preparation of strand-specific RNA-seq libraries using the TruSeq Stranded Total RNA with Ribo-Zero Gold kit (cat# RS-122-2393, Illumina, San Diego, CA, USA). According to manufacturer’s protocol, total RNA was depleted of mitochondrial and cytoplasmic ribosomal RNA followed by fragmentation and random priming to synthesize cDNA fragments. Barcoded sequencing adapters were ligated to cDNA inserts and enriched using PCR to create the final cDNA libraries. Qualitative and quantitative assessment of libraries was performed using a Fragment Analyzer (Advanced Analytical Technologies, Ankeny, IA, USA). Quantified, barcoded libraries were normalized and mixed at equimolar concentrations into a multiplexed sequencing library.

#### RNA-sequencing and data processing

Pooled libraries were sequenced on a HiSeq 4000 sequencer (Illumina, San Diego, CA, USA) to a depth of 30 million reads per sample using a paired-end 100 base pair run configuration. Four samples were sequenced in each pool to correct for potential batch effect and longitudinal samples from the same participants were mixed in the same pool. Sequencing data were demultiplexed and converted into fastq files using Illumina’s bcl2fastq conversion software (v2.20). Quality and adapter trimming along with filtering of rRNA reads was performed using BBDuk (v37.22). The decontaminated reads were mapped to personal genomes using STAR aligner (v2.5.1b) by modifying the GRCh38 reference genome at variant sites called for each participant through exome sequencing. Gene quantification was performed using the tool htseq-count from the Python package HTSeq (v0.9.1). The GENCODE v28 annotation was used to define genomic features where each gene is considered as the union of all its exons. After normalization of read counts to the sequencing depth in each sample, genes with an average expression below 10 were discarded. Missing values were imputed using the k-nearest neighbors’ method (‘impute.knn’ function) in the R package ‘impute’ (v1.52.0). Two datasets were generated, one containing read counts normalized to the sequencing depth in each sample (original), and another that was further processed by applying the variance-stabilizing transformation (VST) in R package ‘DESeq2’ (v3.9).

### Untargeted Proteomics from Plasma by Sequential Window Acquisition of all Theoretical (SWATH)-MS

#### Sample preparation and data acquisition

Plasma samples were thawed on ice, prepared and analyzed in a randomized order. Tryptic peptides were generated from 8 µg of undepleted plasma proteins and separated on a NanoLC 425 System (SCIEX). 5 µl/min flow was used with trap-elute setting using a ChromXP C18 trap column 0.5 x 10 mm, 5 µm, 120 Å (SCIEX, cat# 5028898). Tryptic peptides were eluted from a ChromXP C18 column 0.3 x 150 mm, 3 µm, 120 Å (SCIEX Cat# 5022436) using a 43-minute gradient from 4-32% B with 1-hour total run. Mobile phase solvents consisted of 92.9% water, 2% acetonitrile, 5% dimethyl sulfoxide, and 0.1% formic acid (A) and 92.9% acetonitrile, 2% water, 5% dimethyl sulfoxide (DMSO), and 0.1% formic acid (B). MS analysis was performed using SWATH Acquisition on a TripleTOF 6600 System equipped with a DuoSpray Source and 25 μm I.D. electrode (SCIEX). Variable Q1 window SWATH Acquisition methods (100 windows) were built in high sensitivity MS/MS mode with Analyst TF Software (v1.7). A quality control (QC) consisting of an equimolar pool of all the samples in the study was injected at the beginning and end of each batch. Samples were run in two batches and QC data were used to control for batch effect. Longitudinal samples from the same participants were run in the same batch.

#### Data processing

Peak groups from individual runs were statistically scored with pyProphet tool (v2.0.1) and all runs were aligned using TRIC strategy (Rost et al., 2016). A final data matrix was produced with 1% FDR at peptide level and 10% FDR at protein level. Protein abundances were computed as the sum of the three most abundant peptides (top3 method). Batch effect was corrected by applying median-normalization and proteins detected in less than 2/3 of the samples were discarded. Missing values were imputed by drawing from a random distribution of low values in the corresponding sample (Tyanova et al., 2016). Untargeted protein levels were reported as spectral counts.

### Targeted Proteomics from Plasma by Immunoassays

Plasma samples were thawed on ice, prepared and analyzed in a randomized order. Levels of circulating cytokines and growth factors were measured in plasma using a 63-plex Luminex antibody-conjugated bead capture assay (eBiosciences/Affymetrix). Metabolic hormones were measured using MILLIPLEX MAP Human Metabolic Hormone Magnetic Bead Panel - Metabolism Multiplex Assay (HMHEMAG-34K, Millipore). Cardiovascular risk markers in plasma were measured using MILLIPLEX MAP Human Cardiovascular Disease (CVD) Magnetic Bead Panel (1 to 4) - Cardiovascular Disease Multiplex Assay (HCVD1MAG-67K, HCVD2MAG-67K, HCVD3MAG-67K, HCVD4MAG-67K, Millipore). Experiments were performed by the Stanford Human Immune Monitoring Center (HIMC) according to the manufacturer’s recommendations and read using a Luminex 200 instrument with a lower bound of 20 beads per sample per analyte. Custom assay control beads by Radix Biosolutions were added to all wells for Human 63-plex assay. Longitudinal samples from the same participant were analyzed on the same plate. Inter-plate variability was corrected using the median of inter-plate ratios for the four representative samples analyzed in each plate. Raw mean fluorescence intensity (MFI) values were used for the analysis. Missing values (bead count below 20) were imputed using the k-nearest neighbors’ method (‘impute.knn’ function) in the R package ‘impute’ (v1.52.0). Targeted protein levels were reported as MFI.

### Untargeted Metabolomics from Plasma by Liquid Chromatography (LC)-MS

#### Sample preparation and data acquisition

Plasma samples were thawed on ice, prepared and analyzed randomly as previously described (Contrepois et al., 2015; Piening et al., 2018; Zhou et al., 2019). Briefly, metabolites were extracted using 1:1:1 acetone:acetonitrile:methanol, evaporated to dryness under nitrogen and reconstituted in 1:1 methanol:water before analysis. Metabolic extracts were analyzed four times using HILIC and RPLC separation in both positive and negative ionization modes. Data were acquired on a Thermo Q Exactive plus mass spectrometer for HILIC and a Thermo Q Exactive mass spectrometer for RPLC. Both instruments were equipped with a HESI-II probe and operated in full MS scan mode. MS/MS data were acquired on quality control samples (QC) consisting of an equimolar mixture of all samples in the study. HILIC experiments were performed using a ZIC-HILIC column 2.1 x 100 mm, 3.5 μm, 200Å (Merck Millipore) and mobile phase solvents consisting of 10 mM ammonium acetate in 50/50 acetonitrile/water (A) and 10 mM ammonium acetate in 95/5 acetonitrile/water (B). RPLC experiments were performed using a Zorbax SBaq column 2.1 x 50 mm, 1.7 μm, 100Å (Agilent Technologies) and mobile phase solvents consisting of 0.06% acetic acid in water (A) and 0.06% acetic acid in methanol (B). Data quality was ensured by (i) injecting 6 and 12 pool samples to equilibrate the LC-MS system prior to run the sequence for RPLC and HILIC, respectively, (ii) injecting a pool sample every 10 injections to control for signal deviation with time, and (iii) checking mass accuracy, retention time and peak shape of internal standards in each sample.

#### Data processing

Data from each mode were independently analyzed using Progenesis QI software (v2.3) (Nonlinear Dynamics). Metabolic features from blanks and that didn’t show sufficient linearity upon dilution in QC samples (r < 0.6) were discarded. Only metabolic features present in >2/3 of the samples were kept for further analysis. Inter- and intra-batch variation was corrected using the LOESS (locally estimated scatterplot smoothing Local Regression) normalization method on QC injected repetitively along the batches (span = 0.75). Data were acquired in three and two batches for HILIC and RPLC modes, respectively. Missing values were imputed by drawing from a random distribution of low values in the corresponding sample (Tyanova et al., 2016). Data from each mode were merged and metabolites were formally identified by matching fragmentation spectra and retention time to analytical-grade standards when possible or matching experimental MS/MS to fragmentation spectra in publicly available databases. After careful annotation of the metabolite dataset, a total of 728 metabolites were measured and categorized in classes and pathways based on the KEGG database where possible. Some metabolites elute in multiple peaks and are indicated with a number in parenthesis following the metabolite name ordered by elution time. Metabolite abundances were reported as spectral counts.

### Semi-targeted Lipidomics from Plasma using the Lipidyzer Platform

#### Sample preparation

Plasma samples were thawed on ice, prepared and analyzed in a randomized order. Plasma lipids were extracted using a biphasic separation protocol with ice-cold methanol, methyl tert-butyl ether (MTBE) and water (Contrepois et al., 2018). Briefly, 300 μl of methanol spiked-in with internal standards (provided with the Lipidyzer platform) was added to 40 μl of plasma and vortexed for 20 s. Lipids were solubilized by adding 1,000 μl of MTBE and incubated under agitation for 30 min at 4°C. After addition of 250 μl of ice-cold water, the samples were vortexed for 1 min and centrifuged at 14,000 g for 5 min at 20°C. The upper phase containing the lipids was then collected, dried down under nitrogen, reconstituted with 200 μl of methanol and stored at −20°C. The day of the experiment, lipids were dried down under nitrogen and reconstituted with 300 μl of 10 mM ammonium acetate in 9:1 methanol:toluene.

#### Data acquisition and processing

Lipid extracts were analyzed using the Lipidyzer platform that comprises a 5500 QTRAP System equipped with a SelexION differential mobility spectrometry (DMS) interface (SCIEX) and a high flow LC-30AD solvent delivery unit (Shimazdu). A full description of the method is available elsewhere (Contrepois et al., 2018). Briefly, lipid molecular species were identified and quantified using multiple reaction monitoring (MRM) and positive/negative ionization switching. Two acquisition methods were employed covering 10 lipid classes; method 1 had SelexION voltages turned on while method 2 had SelexION voltages turned off. Lipidyzer data were reported by the Lipidomics Workflow Manager (LWM) software which calculates concentrations for each detected lipid as average intensity of the analyte MRM/average intensity of the most structurally similar internal standard (IS) MRM multiplied by its concentration. Lipid abundances were reported as concentrations in nmol/g. Lipids detected in less than 2/3 of the samples were discarded and missing values were imputed by drawing from a random distribution of low values in the corresponding sample (Tyanova et al., 2016). Data quality was ensured by i) tuning the DMS compensation voltages using a set of lipid standards (SCIEX, cat#:5040141) after each cleaning, more than 24 hours of idling or 3 days of consecutive use, ii) performing a quick system suitability test (QSST) (SCIEX, cat#: 50407) before each batch to ensure acceptable limit of detection for each lipid class, and iii) triplicate injection of lipids extracted from a reference plasma sample (SCIEX, cat#: 4386703) at the beginning of the batch. The data were acquired in eight batches. Two datasets were generated, one containing all individual lipid species (original) and a second one containing concatenated TAG information and LPC, LPE, FFA classes were discarded (redundant with metabolomic dataset). In the latter, the signal from individual TAG with the same total number of carbons and unsaturations were summed (TAG_sum).

### 16S Microbiome Sequencing from Stool

Stool samples were collected within 4 months of the day of exercise (112.9 days in average) as part of the NIH integrated Human Microbiome Project 2 study. DNA was first extracted from stool in line with the Human Microbiome Project’s (HMP) Core Sampling Protocol A (hmpdacc.org) and then sequenced using 2 x 300 bp paired-end sequencing (Illumina MiSeq). Raw sequences were processed using Illumina’s software, assigned to Operational Taxonomic Units (OTU) by Usearch against GreenGenes database (May 2013 version) and final taxonomic assignment was performed using RDP-classifier (Zhou et al., 2019).

### Clinical laboratory tests

Clinical laboratory tests were performed within 2 months of the day of exercise (54.2 days in average) as part of the NIH integrated Human Microbiome Project 2 study.

### Data Analysis

When not specified otherwise, the transcriptomic dataset used was transformed with variance-stabilization (VST), the lipid dataset contained individual TAG species (original), and all missing values were imputed.

#### Fuzzy c-mean clustering

Fuzzy c-mean clustering was performed using the R package ‘Mfuzz’ (v2.20.0) after log2-transformation and Z-score scaling of the data. We calculated the minimum centroid distance for a range of cluster numbers and the optimal number was chosen using the ‘elbow’ method.

#### t-distributed stochastic neighbor embedding (tSNE) dimensionality reduction

tSNE scatterplots were generated after log2-transformation and Z-score scaling of the data using the R package ‘Rtsne (v0.15) with the following parameters: perplexity = 5, theta = 0.05.

#### Inter-individual variability

Coefficients of variation (CV) were calculated for each analyte on non-imputed datasets to avoid any potential bias. CV = standard deviation/mean*100. The transcriptomic dataset used was normalized to sequencing depth (original). Inter-individual variability in response to exercise was determined as the median CV for each analyte across all time points post-exercise relative to baseline. Technical variability for each assay was determined by calculating the CV across all quality control samples in the study.

#### Linear regression analysis to find analytes that change in response to exercise

Linear models were applied on log2-transformed data. Lipid dataset used was the concatenated TAG version (TAG_sum). The model corrected for personal baseline, age, sex, body mass index, ethnicity/race and batch information. Linear models were computed in R using the lm base function. *P* values were corrected for multiple hypothesis using the Benjamini-Hochberg (BH) method and analytes with FDR below 0.05 were considered significant. ANOVA testing was performed on significant analytes to compute *P* values at each time point post-exercise. These *P* values were also corrected using BH method and analytes with an FDR below 0.05 were considered significant.

#### Pathway/chemical class enrichment analysis

We used the Ingenuity pathway analysis (IPA) platform to search for enriched pathways using differentially expressed genes in PBMCs and circulating proteins. All the detected transcripts or proteins were used as a background. Significance of pathways was determined by the hypergeometric test (one-sided). For metabolites and complex lipids, enrichment was calculated using the Kolmogorov–Smirnov method (Barupal and Fiehn, 2017) in R with pathway/chemical class annotation of all detected metabolites/lipids. This method uses distribution of *P* values and does not rely on the size of background databases. *P* values were corrected for multiple hypothesis using the Benjamini-Hochberg method and pathways with FDR below 0.05 (transcripts, proteins) and 0.20 (metabolites, complex lipids) were considered significant. Pathway directions were calculated as follows: 1) median of fold change values relative to baseline for significant molecules in the pathway, 2) median of beta coefficients for significant molecules in the pathway, 3) median of max (if up) or min (if down) fold change values relative to baseline for significant molecules in the pathway. Beta coefficients were provided by the regression model (see below).

#### Correlation network analysis

Pairwise Spearman’s rank correlations were calculated using the R package ‘Hmisc’ (v4.1-1) and weighted, undirected networks were plotted with ‘igraph’ (v1.2.1). Correlations with Bonferroni adjusted *P* values below 0.05 were included and displayed via the fruchterman-reingold method. Only the main network was plotted. Nodes were color-coded by assay and their size represent the maximum (clusters 1 and 2) or minimum (clusters 3 and 4) median fold change in response to exercise across participants in the cohort.

#### Differential allele-specific expression (ASE) with exercise

Read mapping bias was removed by following the WASP pipeline. The *GATK* tool *ASEReadCounter* was used to count allele specific reads at exonic heterozygous sites. For each individual, only SNPs supported by at least 20 reads in each time point and bi-allelic (0.1 ≤ allelic ratio ≤ 0.9) in at least one time-point were considered. Only SNP shared by at least 10 heterozygous individuals were considered. We tested for differences in ASE with time using the beta-binomial generalized linear mixed models implemented in EAGLE (v2.0) (Knowles et al., 2017). In short, for each exonic SNP, we model the alternative allele count for heterozygous individual *i*, at time *t* using a Beta-Binomial distribution, i.e. yit ∼ BB(nit, σ(φi βt + ui), c), where nit is the total number of reads for *i* in *t*, σ() is the logistic function, φ*i* represents the phase between the causal cis-SNP and the exonic SNP and is treated as a latent variable taking values in {−1,+1}. We learn a prior π = P(φi = +1) across all individuals and marginalize (sum) over the possible values of π. βt is the effect of time on the reference allele proportion. ui ∼ N(0,v) is a per individual, per exonic SNP random effect. We use variational Bayes EM to approximately integrate over all ui while optimizing with respect to the other parameters. Last, c is the concentration parameter which we learn per exonic SNP using maximum a-posteriori probability estimation with a Gamma(1.0001, 10E−4) prior. We use a likelihood ratio test to test the global hypothesis of no differences in ASE between baseline and any time point. We corrected for multiple testing across SNPs using the BH procedure (FDR < 0.05).

#### Linear regression to find analytes associated with peak VO_2_

Linear regressions were conducted at each time point using the lm base function in R and corrected for age, sex, body mass index, ethnicity/race and batch information. *P* values were corrected using BH method and analytes with a *P* value below 0.05 were considered significant. The lipid dataset used was TAG_sum.

#### CPX parameter prediction models

Prediction models were trained using baseline multi-omic measurements (pre-exercise). Clinical laboratory results and gut microbiome data were from the iHMP dataset at the closest healthy visit to the exercise day. No clinical and gut microbiome data were available for participant ZJBOZ2X, hence this participant was discarded from the analysis. Prediction modeling was performed by first selecting predictive analytes via Bayesian networks and then build the model using ridge regression (Schussler-Fiorenza Rose et al., 2019).

##### Feature selection

All data (demographics, clinical, complex lipids, metabolites, targeted proteins, proteins, transcripts) except gut microbiome percent data were log2-transformed. All features were standardized to zero mean with unit variance. Output CPX parameters were not transformed or scaled. Identification of predictive features was performed using the Max-Min Parents and Child algorithm (MMPC) in the R package MXM (v1.4.1). Feature selection was performed using leave-one-out cross-validation, where 34 training sets were constructed and each training set excludes the data from a different participant. MMPC algorithm was run on each training set and predictive features were selected if they were used in ≥ 20% of the models generated from the training sets.

##### Ridge regression modeling

We performed leave-one-out cross-validation to maximize available training data. For each training set, we optimize the hyperparameter by performing a grid search and selecting the model that minimizes test error. The predicted output value is the value from the cross-validation iteration in which that output data point and its associated features are excluded from the training set. We use these predicted values to calculate mean square error (MSE) and R^2^. The value of the hyperparameter used was the average of the hyperparameters which minimized test error during cross-validation.

#### Linear mixed models to find analytes responding differently to exercise in insulin resistant participants

Linear mixed models were conducted at each time point using the ‘lme4’ package (v1.1-21) in R and corrected for personal baseline, age, gender, BMI, race/ethnicity and batch information. ‘lmerTest’ package in R (v3.1-0) was used to compute *P* values at each time point post-exercise. *P* values were corrected using BH method and analytes with a *P* value below 0.05 were considered significant. The lipid dataset used was TAG_sum.

### Multi-omic outlier analysis

Outlier analysis was performed on non-imputed datasets to avoid any potential bias. Multi-omic data were log2-transformed and all features were standardized to zero mean with unit variance. Absolute levels and delta values relative to baseline were used for the ‘baseline’ and ‘in response to exercise’ analyses. *P* values were calculated assuming a normal distribution and were corrected for multiple hypothesis using the Benjamini-Hochberg procedure. Analytes with a *P* value below 0.05 at baseline were considered outliers. For outliers in response to exercise, two significant outliers across the four time points (2, 15, 30 min and 1 hour time points) were required.

---

**Figure S1.**
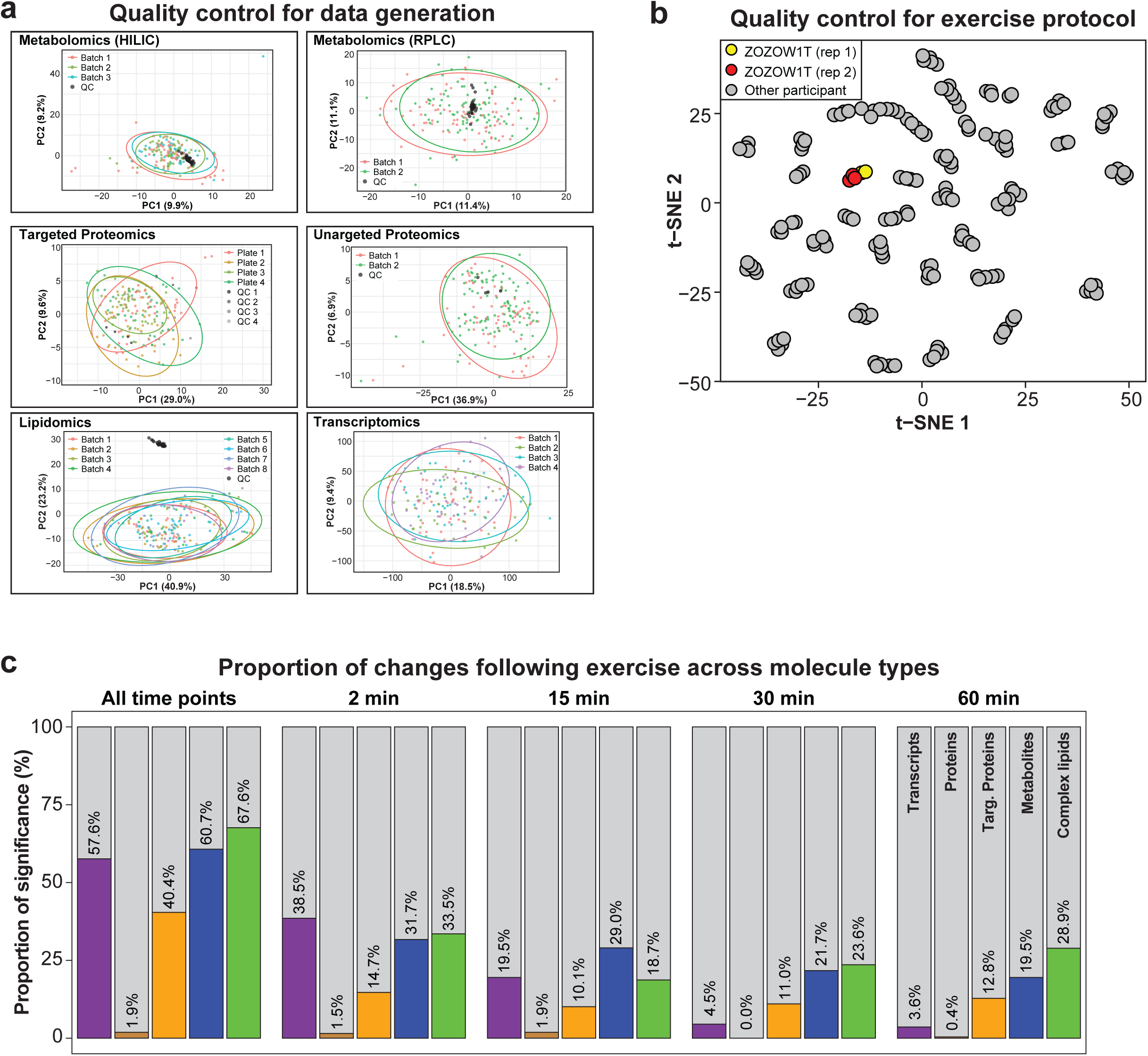
Quality controls for data generation and exercise protocol as well as molecular changes in response to exercise. (**a**) Principal component analysis across omic assays. Each dot represents a sample colored by batch information. Black and grey dots are quality control samples (QCs). For metabolomics and untargeted proteomics, QCs consist in an equimolar mix of all the samples in the study. For lipidomics, QCs are lipid extracts from a reference plasma sample. QC1-4 in targeted proteomic experiments represent individual samples in the study analyzed in each plate. Transcriptomic experiment didn’t contain QC. (**b**) 2D visualization of all multi-omic analytes using t-distributed stochastic neighbor embedding (tSNE) technique. Each dot represents a single sample colored by participants. (**c**) Bar graph representing the proportion of analyte across molecule types significantly changing in response to exercise (FDR < 0.05).

**Figure S2.**
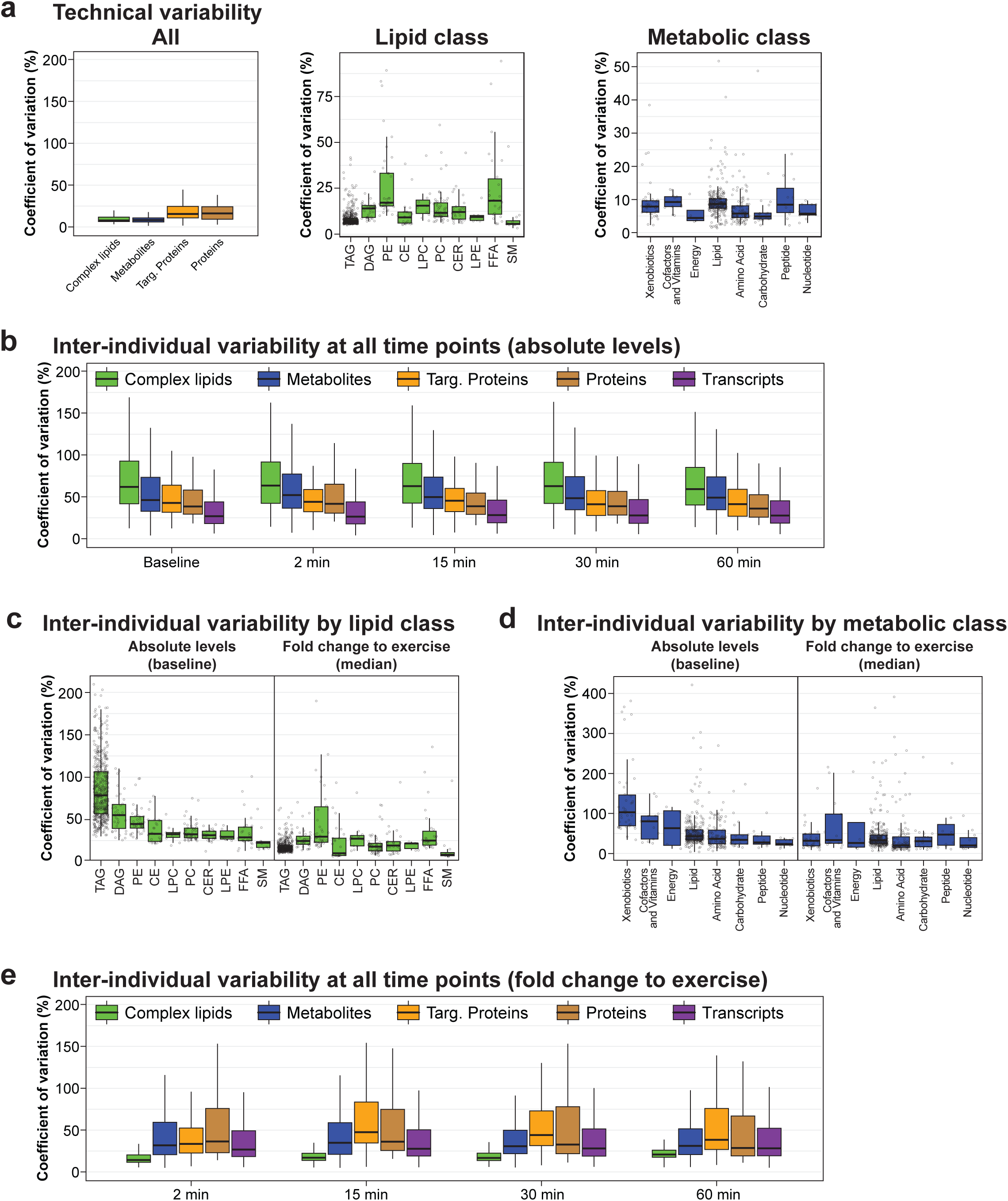
Inter-individual variability. (**a**) Technical variability for complex lipids, metabolites and proteins using quality control samples as described in **Figure S1a**. (**b**) Inter-individual variability at all time points using absolute levels across molecule types. Inter-individual variability by lipid class (**c**) and metabolic class (**d**). (**e**) Inter-individual variability at all time points using analyte levels relative to baseline.

**Figure S3.**
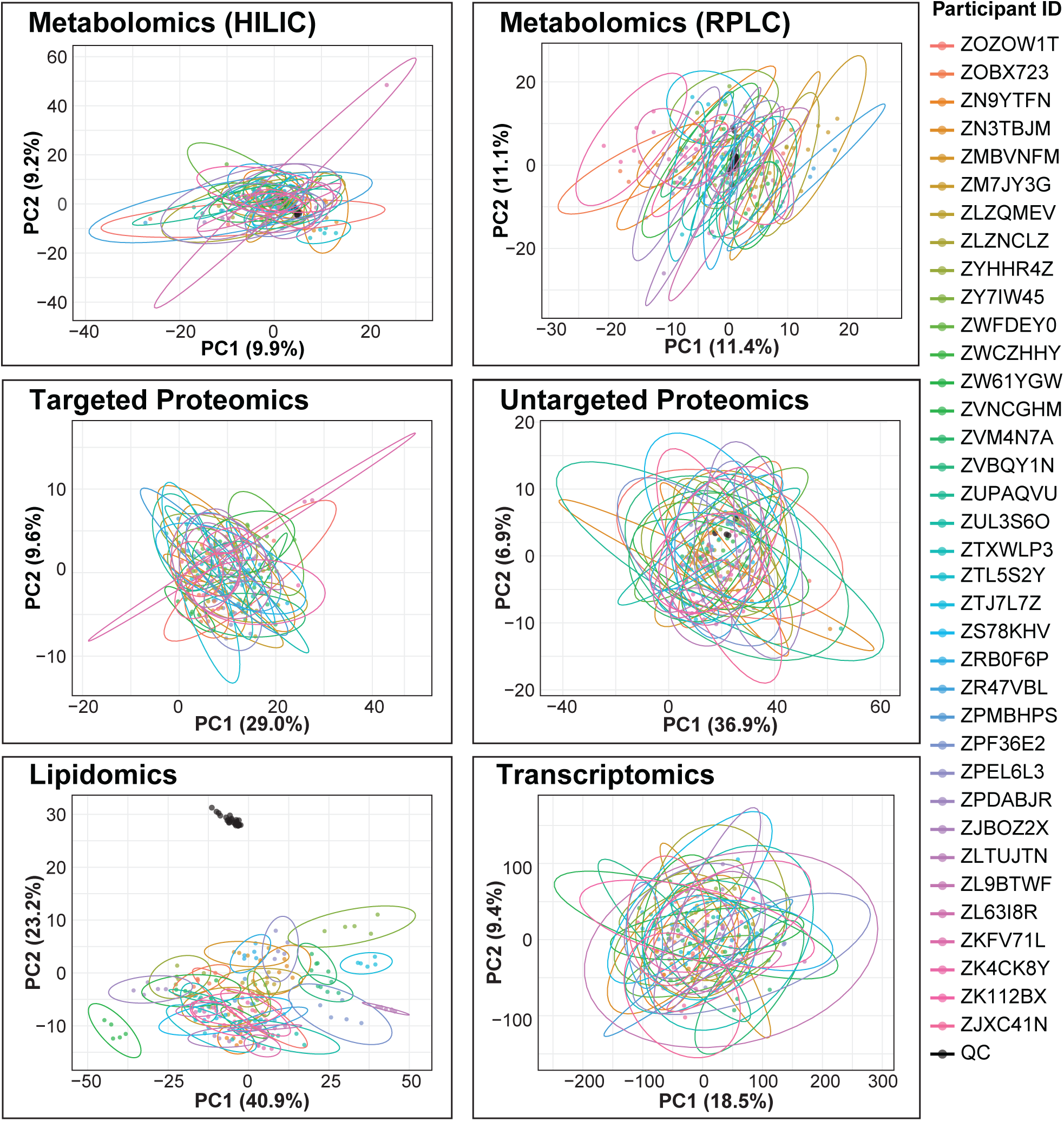
Individuality across molecule types. Principal component analysis across omic assays. Each dot represents a sample colored by participants.

**Figure S4.**
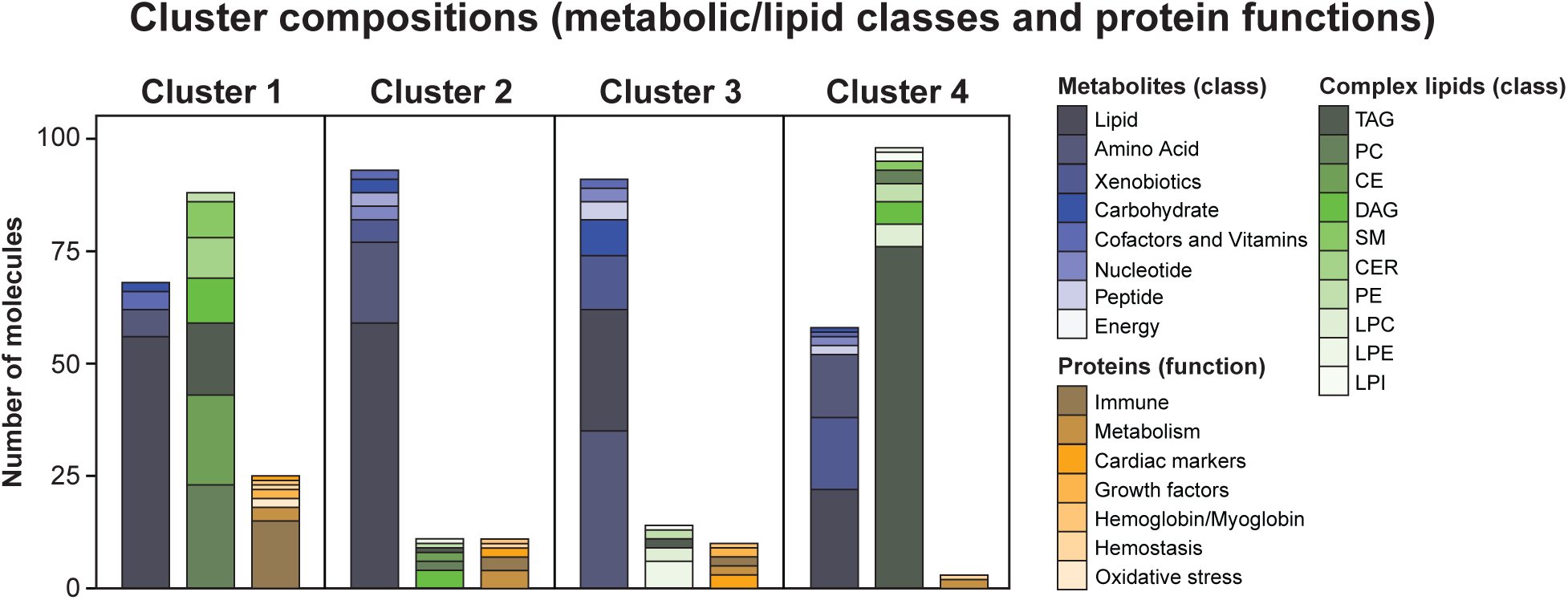
Cluster compositions in metabolite and complex lipid classes as well as protein functions.

**Figure S5.**
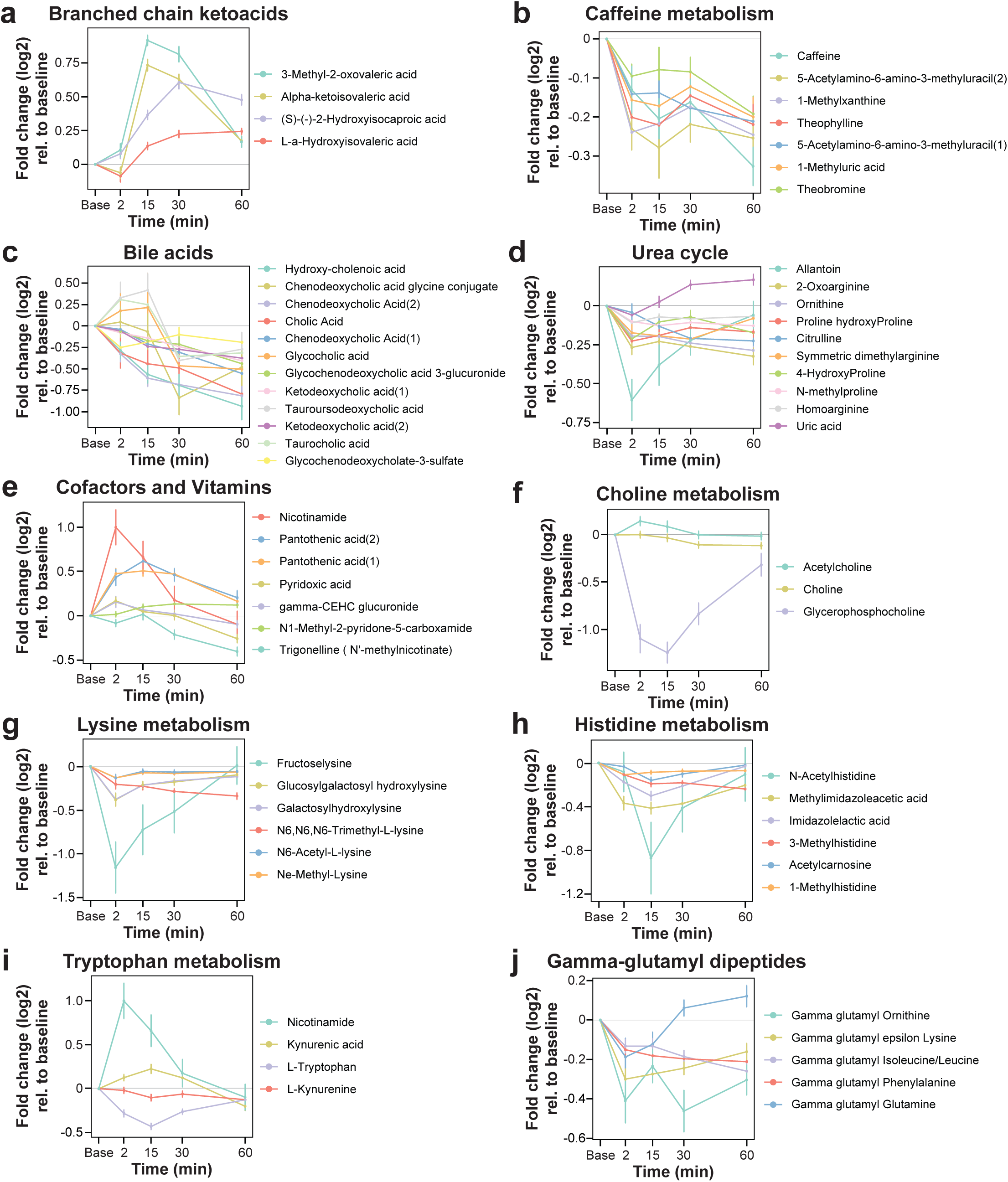
Longitudinal trajectories of selected metabolic pathways in response to exercise. (**a**) branched-chain ketoacids, (**b**) caffeine metabolism, (**c**) bile acids, (**d**) urea cycle, (**e**) cofactors and vitamins, (**f**) choline metabolism, (**g**) lysine metabolism, (**h**) histidine metabolism, (**i**) tryptophan metabolism and (**j**) gamma-glutamyl dipeptides.

**Figure S6.**
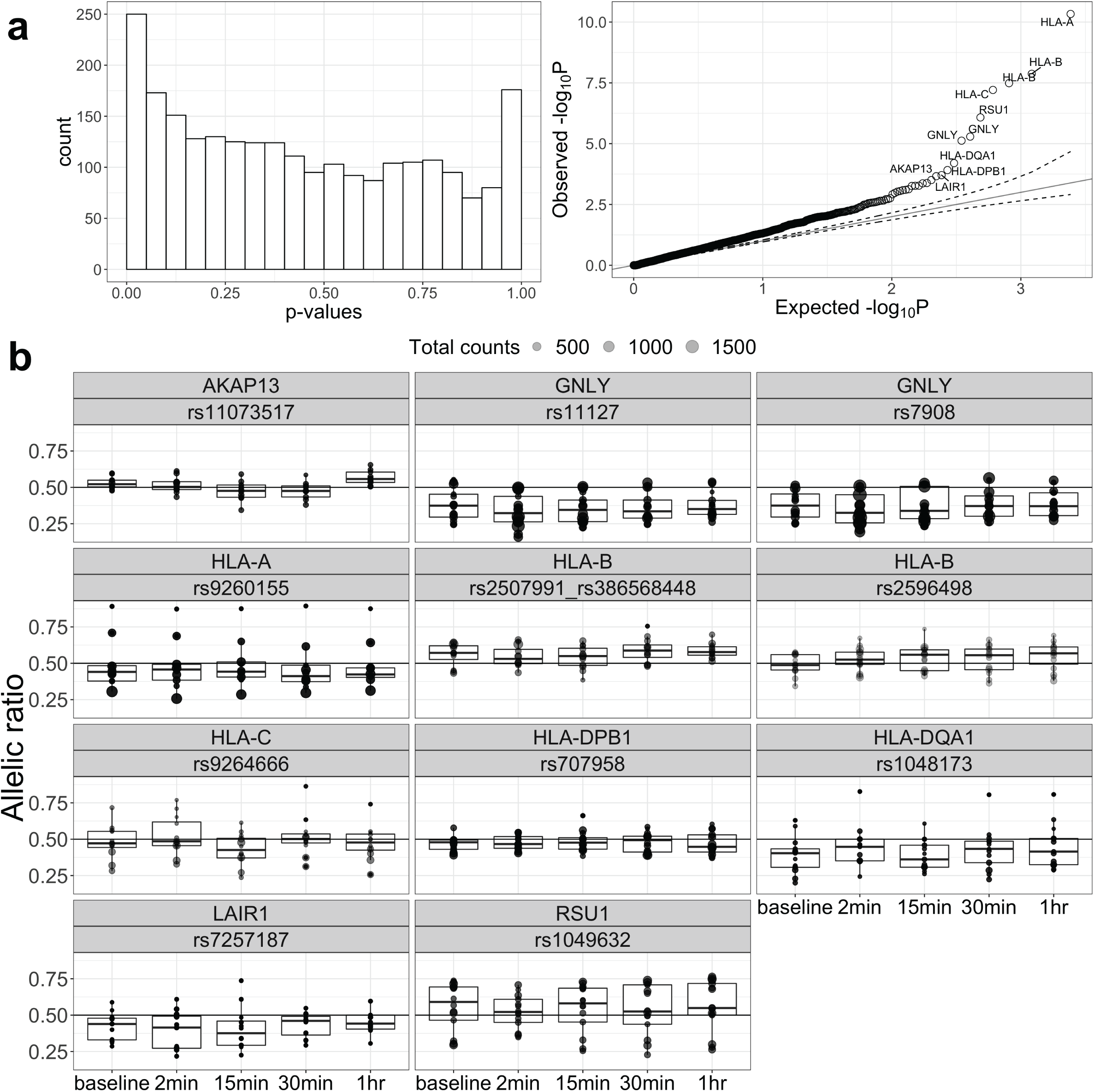
Population-level differential allele-specific expression (ASE) by time. (**a**) Histogram and QQ plot of likelihood-ratio test (LRT) p-values for differential ASE with time analyses. (**b**) Distribution of allelic ratios across time for all SNPs with significant differential ASE by time effects.

**Figure S7.**
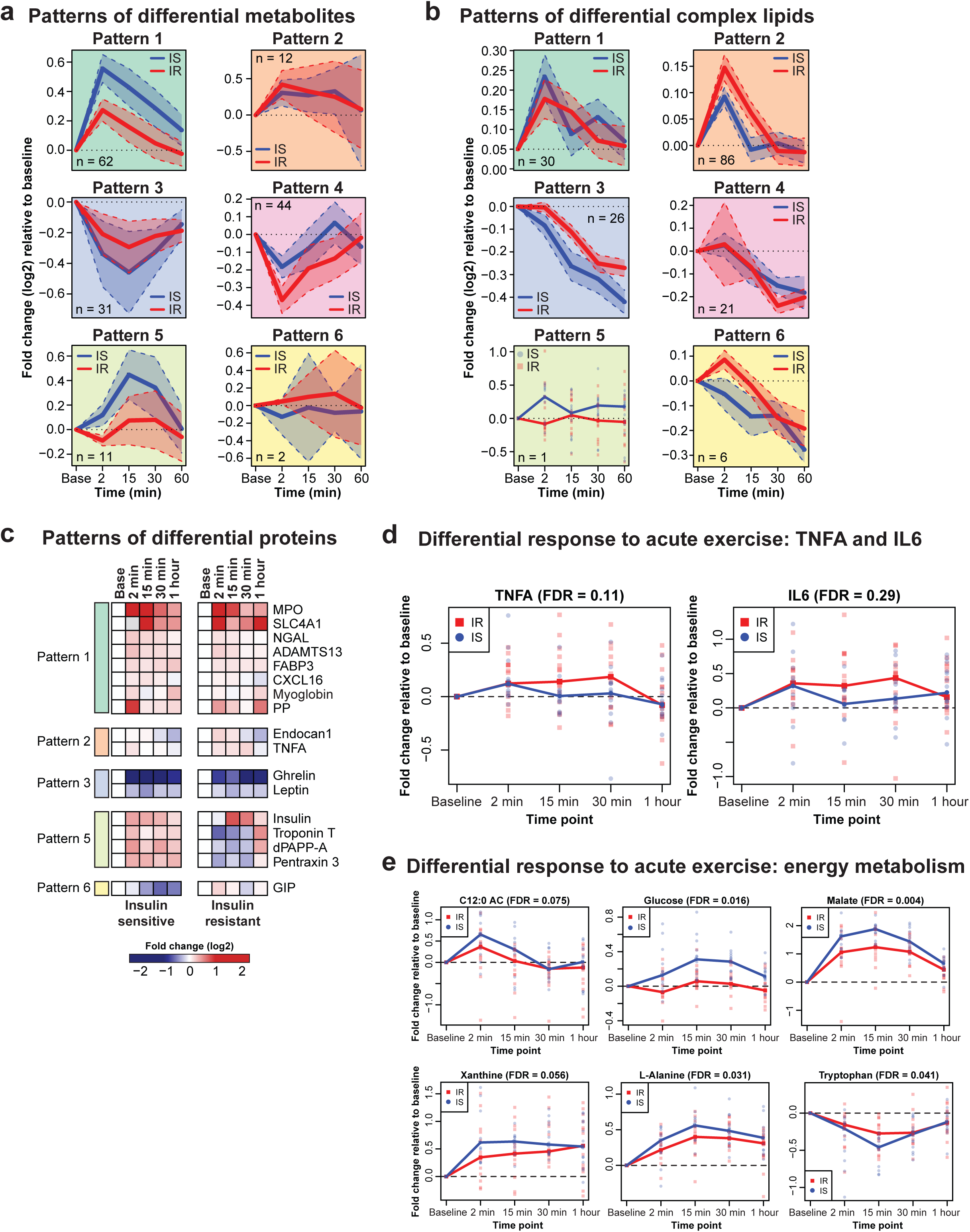
Differential response to acute exercise in insulin resistant participants. Patterns of differential metabolites (**a**) and complex lipids (**b**) in IS and IR participants. Linear mixed models correcting for personal baseline, age, sex, body mass index and race/ethnicity were employed. *P* values were corrected for multiple hypothesis using the Benjamini-Hochberg method. Metabolites and complex lipids were considered significant with FDR below 0.10 and 0.20, respectively. The solid line represents the mean and the dashed line represents the 95% confidence interval. (**c**) Heatmap of significant proteins as determined by the linear mixed model (FDR < 0.20) representing the median log2 fold change relative to baseline in the cohort. Proteins were grouped in patterns. Differential response to exercise for (**d**) TNF-α and IL-6 as well as key metabolites (**e**).

**Figure S8.**
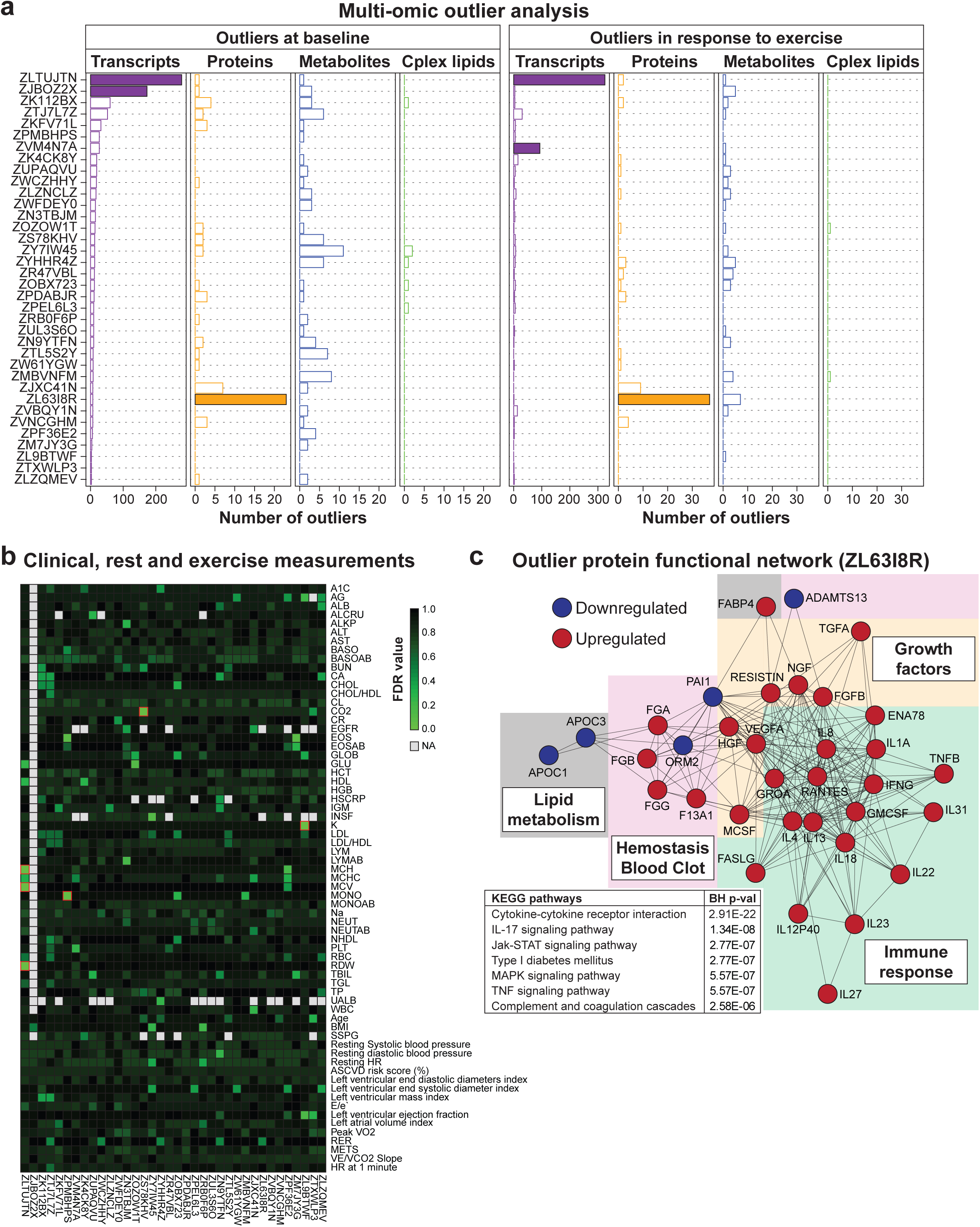
Multi-omic outlier analysis. (**a**) Number of outliers across omes per participant at baseline and in response to exercise (FDR < 0.05). *P* values were corrected for multiple hypothesis using the Benjamini-Hochberg (BH) method. Participants with significantly different number of outliers in comparison to the population (FDR < 0.05) are represented with a solid bar. (**b**) Heatmap of clinical laboratory results as well as key rest and stress parameters representing the BH-corrected *P* values for each measure in each participant. Outlier measures (FDR < 0.05) are outlined in red. (**c**) Functional association network of outlier proteins (FDR < 0.05) in response to exercise in individual ZL63I8R. This analysis was performed using the web tool STRING. Edges correspond to known, predicted or other interactions. Proteins in blue are downregulated and proteins in red are upregulated.

## Supplementary tables

Table S1. Listing of immune proteins, cardiovascular biomarkers and metabolic proteins detected from the targeted panel

Table S2. Listing of all plasma and cellular analytes detected in the study

Table S3. Participants baseline demographics, echocardiographic and cardiopulmonary exercise characteristics

Table S4. Analytes significantly changing in response to acute exercise and their cluster information (simple linear models)

Table S5. Inter-individual variability using absolute levels

Table S6. Pathway enrichment analysis using genes with a CV > 100%

Table S7. Inter-individual variability in response to exercise using fold change to exercise

Table S8. Lipid class and metabolic pathways changing in response to exercise

Table S9. PBMC RNA pathways changing in response to exercise

Table S10. Molecular associations with Peak VO2

Table S11. Lipid classes and metabolic pathways associated with peak VO2

Table S12. RNA pathways associated with Peak VO2

Table S13. Protein pathways associated with Peak VO2

Table S14. Clinical laboratory results (closest to exercise date)

Table S15. Gut microbiome data (closest to exercise date)

Table S16. Peak VO2 prediction models

Table S17. VE/VCO2 slope prediction models

Table S18. RER prediction models

Table S19. Molecules responding differentially to acute exercise in IR and IS participants (linear mixed models)

Table S20. Differential PBMC pathways in IR participants

Table S21. Differential lipid classes and metabolic pathways in IR participants

Table S22. Pathway analysis using outlier transcripts

